# Denisovan introgression left differential selection regimes in Humans and Neanderthals on the *SLC30A9* gene

**DOI:** 10.64898/2026.09.23.753770

**Authors:** Jorge Garcia-Calleja, Cesar A Fortes-Lima, Moisès Coll Macià, Oscar Lao, Elena Bosch

## Abstract

Signals of positive selection around the *SLC30A9* gene have been reported in human populations outside Africa. Selection likely acted on a highly differentiated single-nucleotide polymorphism, rs1047626, leading to a non-synonymous substitution in the encoded zinc transporter. Because of the striking similarity between the putatively selected *SLC30A9* haplotype observed in several current human populations and the Denisovan individual, previous work has proposed adaptive introgression. Yet alternative explanations, including ancient human variation, and the precise archaic source —Neanderthal or Denisovan— remained unresolved. Considering the potentially complex evolution of *SLC30A9*, we applied Approximate Bayesian Computation (ABC) algorithms coupled to machine learning to investigate the most plausible evolutionary origin of this substitution. After modelling different evolutionary scenarios with forward-in-time simulations, our results highlight that the most probable scenario is a Denisovan origin of the rs1047626 polymorphism. However, the allele likely introgressed into Neanderthals first and was then passed into non-African modern humans. Moreover, the derived allele frequency for rs1047626 across several African populations is consistent with back-to-Africa migrations. Finally, our ABC analyses indicate strong positive selection in East Asian populations and other out-of-Africa populations, whereas in Neanderthal populations, the selection coefficient was probably neutral or slightly deleterious.

## INTRODUCTION

Past gene flow between anatomically modern humans (AMH) and extinct Denisovan and Neanderthal hominins has left a noticeable genetic legacy in present-day non-African genomes, amounting between 2 and 6% of their total DNA^1–3^. In some cases, introduced variants conferred selective advantages to admixed individuals, facilitating adaptation to the novel environments encountered during the expansion out-of-Africa. Well-characterized examples include traits related to lighter skin pigmentation, cold tolerance, high-altitude adaptation, and immune responses^4–9^.

Previous studies^10^ have reported *SLC30A9* as a candidate of archaic adaptive introgression. This gene encodes the zinc transporter ZnT9, which plays a key role in intracellular zinc homeostasis—particularly within mitochondria and the endoplasmic reticulum—and is therefore essential for cellular metabolism and viability. Multiple genome-wide selection scans^11–13^ have identified strong signatures of positive selection at this locus in non-African populations, including elevated population differentiation and extended haplotype structure. These signals are largely driven by a highly differentiated non-synonymous variant, rs1047626, which results in a methionine-to-valine substitution at position 50 of ZnT9. This substitution flags two major haplotypes segregating worldwide: an ancestral haplotype prevalent in Africa, and a derived haplotype that reaches high frequency or near fixation across populations of Eurasia and Oceania, which shows high affinity to archaic genomes, particularly Denisovans. Moreover, the derived rs1047626 allele has been experimentally shown to modify intracellular zinc handling in mitochondria and the endoplasmic reticulum^10^, speculated to have an effect on cold adaptation.

Despite this evidence, the detailed evolutionary history and the archaic source of the *SLC30A9* haplotype remain unresolved. First, the derived rs1047626 allele and Denisovan-like haplotypes in modern humans are not restricted to non-African populations but are also present at low to moderate frequencies in certain African groups. This observation challenges a simple model of Denisovan introgression, as primarily sub-Saharan African populations have traditionally been considered largely unadmixed with archaic hominins. Second, the high frequency of the derived haplotype observed in European populations is difficult to reconcile with current models of Denisovan admixture, which exclude direct contact between the two^14,15^.

Here, we address these questions by integrating worldwide allele frequency data—including dense sampling across African populations—with haplotype analyses of present-day and ancient human genomes spanning key periods of human–archaic interaction. To investigate the presence of the archaic haplotype in modern human populations, we developed a novel algorithm based on IBD sharing. We then explicitly test multiple evolutionary models for the origin and spread of the Denisovan-like haplotype using a simulation-based inference framework that combines forward-in-time modelling, Approximate Bayesian Computation (ABC), and machine learning. This framework allows us to disentangle competing hypotheses—including multi-step archaic introgression scenarios and ancient standing variation—and to jointly estimate the timing and strength of selection shaping variation at *SLC30A9* in modern human populations.

## RESULTS

### Geographical distribution of the SLC30A9 methionine-to-valine substitution

We compiled allele frequencies of the rs1047626 derived allele (G) across diverse modern-day human populations around the globe (Supplementary Table S1). As previously reported^10^, the derived allele is predominantly found in frequencies above 50% in most non sub-Saharan African groups, being almost fixed in all East Asian, Oceanian, and Native American population groups (Figure 1). The derived allele is present at low to moderate frequencies in Middle Eastern populations (range: 25%-82%)^16–18^, whereas in North African populations it ranges from 50%-57% in Berber-speaking populations from Morocco^19^, to 62%-73% in Egyptians from Egypt^18,20^, and to 79% in Egyptian migrants residing in Sudan known as Copts^21^. By contrast, most sub-Saharan African populations have very low frequencies or a complete absence of the derived variant (e.g., the derived variant is absent in 57 populations and 106 populations display frequencies below 5%), with a few of them presenting moderate and high frequencies for the derived allele (see detailed pie-charts and surface maps across Africa in Supplementary Figure S1).

**Figure 1.**
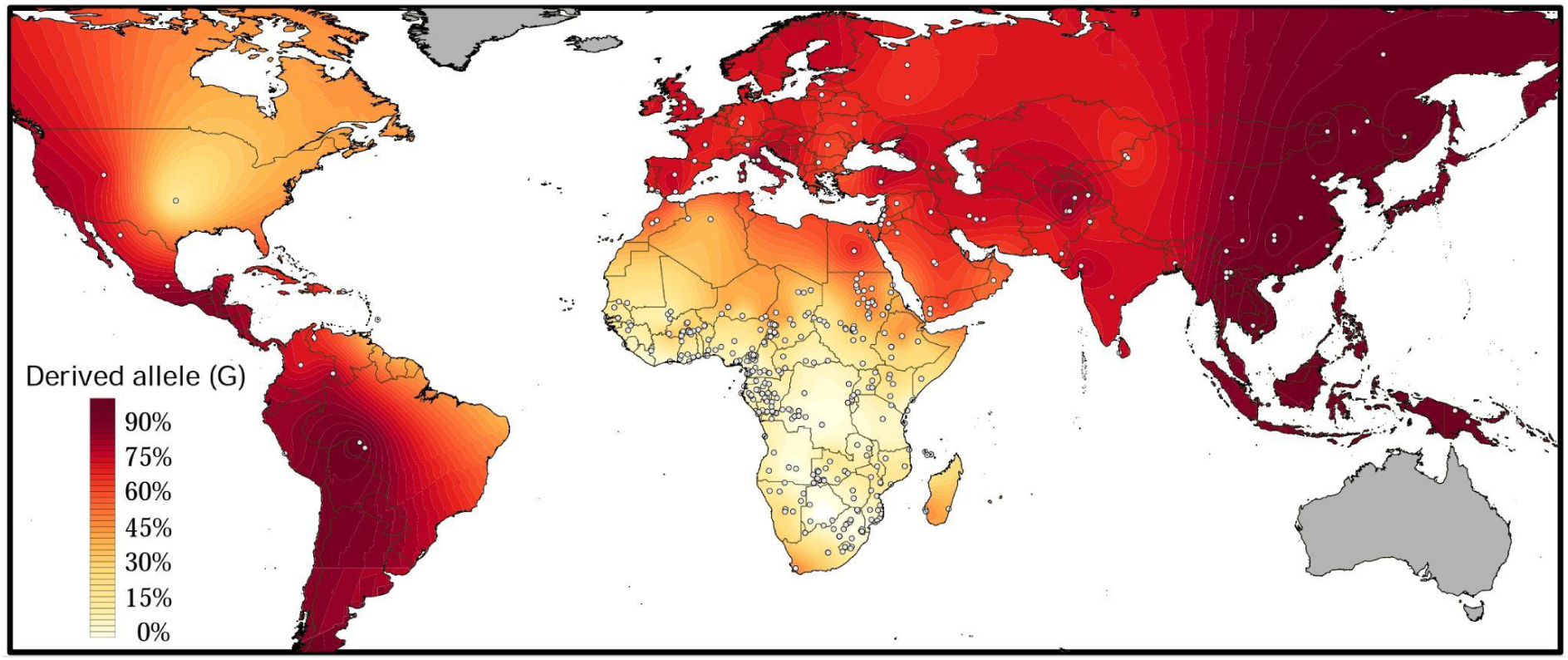
Surface map of the derived allele frequency at rs1047626 locus across worldwide populations using the Kriging spatial interpolation method^35^. Each black dot represents a sampled human population (see complete list and references in Supplementary Table S1). Lighter colours denote low frequencies for the derived variant, while darker colours denote high frequencies. In grey, continental regions for which there are not enough data and no interpolation was considered.

The presence of the methionine-to-valine substitution in sub-Saharan African populations challenges the recognition of *SLC30A9* as introgressed from archaic hominins, since archaic introgression is usually restricted to alleles absent in these populations. However, events of back-to-Africa migrations^22–24^ and several subsequent historical migrations contributing additional genetic components from Western Eurasia could also explain the presence of the derived allele in this continent^2^. Supporting this hypothesis, we find a strong robust linear correlation (r = 0.5945, p-value = 1.8e-9) between the derived allele frequency and the proportion of Eurasian ancestry inferred at K = 4 using unsupervised ADMIXTURE analysis^25^ across African populations (Supplementary Figure S2, Supplementary Table S2).

More specifically, across the Sahel belt, populations known to present non-African ancestry show the highest frequencies of the derived allele, such as Niger-Congo Fulani speakers and Afro-Asiatic Arabic speakers (Supplementary Table S1). In particular, the derived allele frequency of rs1047626 ranges from 21% to 47% in Fulani populations from Chad, Niger, Mali, and Burkina-Faso^26–29^. The derived allele is present at high frequencies in populations that have experienced recent admixture such as the South African Coloured individuals (range: 25%-50%)^25,30^, and populations from the Comoros Islands (range: 12-38%)^31^ and Madagascar (range: 40-50%)^32^. On the contrary, none of the most divergent lineages within Africa, such as the southern and eastern hunter-gatherers (e.g., San and Mbuti, respectively), seem to present the derived allele, or they do so at frequencies below 15%, as observed in the western hunter-gatherers (e.g., 3% in Biaka and 7% in Baka^33,34^).

Altogether, we observe a high frequency of the rs1047626 introgressed derived allele around the globe. In Africa, the modest frequency of the allele can be largely explained by back-to-Africa gene flow that likely introduced the allele in the continent in more recent times. Likewise, populations from the Middle East and the Americas with African ancestry have lower frequencies.

### Haplotype structure of the SLC30A9 region

To further investigate the history and haplotype structure of the *SLC30A9* gene, we examined genotypes surrounding the rs1047626 variant in archaic, ancient, and present-day human samples from the Allen Ancient DNA Resource (ADDR) repository^36^ (Figure 2A, Supplementary Figure S3).

**Figure 2.**
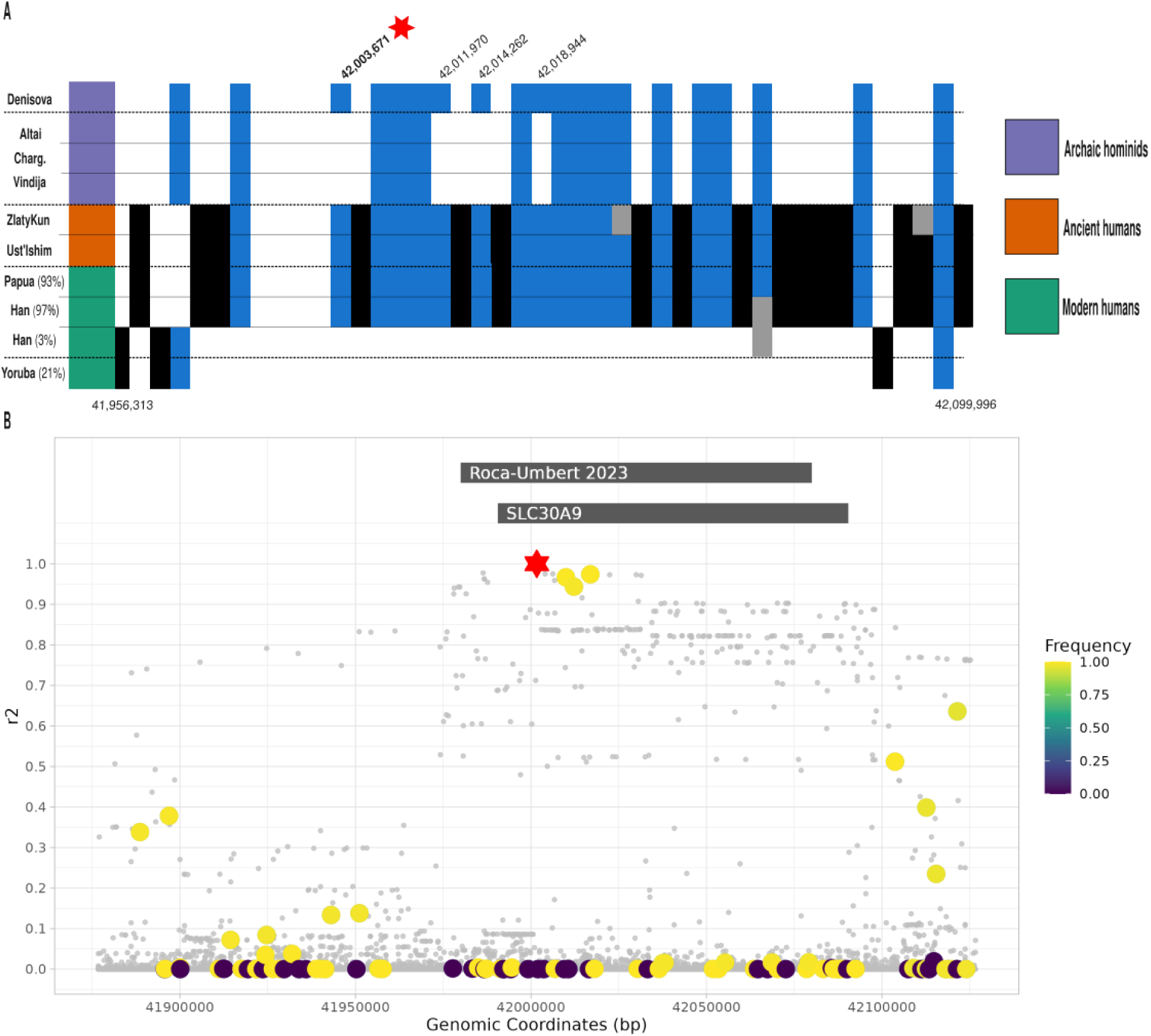
Haplotype structure and linkage disequilibrium of the *SLC30A9* locus in archaic and modern humans. **A)** Schematic representation of the haplotype structure around the *SLC30A9* gene (chr4:41975811-42046424, GRCh37 coordinates) selecting the most representative haplotype for each present-day human population, except for Han Chinese where we show both haplotypes. In blue, alternative alleles shared with Denisovans; in black, alternative alleles not present in Denisovan; in white, reference alleles, and in grey, missing or not-called alleles. Present-day human genomes include Yoruban (1000 Genomes Project), Papuan and Han Chinese (HGDP^42,43^). In parenthesis, the proportion of the haplotype in the population. See Supplementary Figure S3 to see the full population haplotype structure. Archaic hominins represented are Altai Neanderthal (Altai), Altai Denisovan (Denisova), Vindija Neanderthal (Vindija), and Chagryskaya Neanderthal (Charg). Ancient modern humans predating 40,000 years ago include Ust’-Ishim and Zlatý kůň. The Denisovan polymorphisms privately shared with out-of-Africa populations not present in Neanderthals are labelled at the top. The rs1047626 polymorphism corresponds to position 42,003,671, highlighted by a red star. **B)** Linkage disequilibrium (LD) values (diploid r2) between the rs1047626 and variants in a 250 kb region (GRCh38 coordinates). Colourful dots show informative variants that are called in archaic genomes and in which Denisova differs from Neanderthals. The colour scale shows the frequency of the Denisova-like allele in modern human individuals that are homozygous derived for the rs1047626 loci. Strips show the *SLC30A9* location and the high-LD region annotated in Roca-Umbert 2023.

Across a ∼143 kb region encompassing 47 SNPs, archaic sequences are nearly identical, with differences limited to rs1047626 (chromosome 4, 42,003,671, GRCh37)—where the Denisovan genome carries the derived “G” allele and all Neanderthal genomes carry the ancestral “A” allele—and three additional positions (42,011,970, 42,014,262, and 42,018,632; rs2581441, rs2581430, and rs1848180, respectively). Modern Yoruba individuals lacking the derived rs1047626 allele –as proxies of the most common haplotype in Africa– form an outgroup to the archaic clade in this region. In contrast, both ancient and present-day human individuals carrying the derived allele show greater similarity to archaic sequences and share the Denisovan state at all three additional sites. This pattern is consistent with previous observations^10^, supporting the presence of a Denisovan-like haplotype associated with the derived allele. Notably, the apparent clarity of the haplotype structure in Figure 2A likely reflects the use of capture-based genotype data at pre-ascertained sites, which may underestimate the true diversity within the region. Nonetheless, the same conclusions are reached when analyzing whole-genome sequences in modern samples^10^ (Supplementary Figure S4).

To obtain a more individual-level view of the region, we next inferred local ancestry across the genomic region surrounding rs1047626 using jointly called data from the 1000 Genomes Project and the Human Genome Diversity Project (gnomAD-harmonized data)^37^. We applied a supervised haplotype-similarity framework (Methods), assigning ancestry to a set of proxy populations: San (representing modern human ancestry) without the derived allele, three high-coverage Neanderthal genomes, and the Denisovan genome. Among haplotypes carrying the derived rs1047626 allele, we observe that a ∼145 kb region including the rs1047626 variant is predominantly assigned to Denisovan ancestry, in both non-African and a subset of African individuals (Supplementary Figures S5–S8). Flanking regions are predominantly assigned to San ancestry. This Denisovan-like fragment has very similar lengths among all individuals analyzed, even across modern human populations. As a control, haplotypes from Mandenka and CEU individuals that do not carry the derived rs1047626 allele are consistently assigned to San ancestry across the same genomic interval (Supplementary Figures S9–S10), supporting the specificity of the Denisovan signal, as previously suggested^10^.

To better characterize the haplotype structure in modern populations, we examined linkage disequilibrium (LD) patterns using the gnomAD-harmonized data^37^ (Methods). The three SNPs that differ between Denisovans and Neanderthals identified above (Figure 2A) are in strong LD with rs1047626 (r^2^ ≈ 0.95, Figure 2B) and are part of a broader cluster of variants showing high LD (r^2^ > 0.8) across a region of approximately 125 kb. This extended haplotypic structure tags the Denisovan-like introgressed fragment identified by our local ancestry method and coincides with the previously identified haplotype^10^ (Roca-Umbert *et. al.* 2023 strip in Figure 2B). However, no additional variants in strong LD within the region distinguish Denisovan from Neanderthal ancestry, limiting our ability to unambiguously resolve the archaic source based solely on LD patterns and variant matching (Figure 2B, Table 1). Other SNPs that differentiate Denisovan and Neanderthal sequences are located more than 100 kb from rs1047626, exhibit weak LD with this locus, and do not clearly stratify modern human individuals by rs1047626 genotype (Table 1). Therefore, the assignment of Denisovan origin to this region in individuals carrying the rs1047626 derived allele is primarily driven by these four closely linked loci in strong LD. This may reflect incomplete representation of the genetic variation present in the introgressing archaic population in currently sequenced genomes, as several variants in the region show similarly high LD but are not observed in any available archaic sequences.

**Table 1.** Summary statistics of informative sites for the discrimination of archaic origin. Chromosomal positions are shown relative to GRCh38. Diploid r2 values are computed using the gnomAD data. Archaic genome columns display the observed genotype (0/0: homozygous reference; 0/1: heterozygous; 1/1: homozygous alternative; ./.: missing data). The frequency columns report the frequency of the allele in Denisova (0 or 1) for each SNP in 1) individuals that are homozygous derived for the rs1047626 locus (D/D) and 2) individuals that are homozygous for the ancestral allele at the same locus (A/A) in gnomAD data. Only informative variants (Neanderthals are different to Denisova) with r2 > 0.2 to rs1047626 locus are shown in this table.

| CHR | POS | $r^2$ | Vindija | Chagyrskaya | Altai | Denisova | freq (D/D) | freq (A/A) |
| --- | --- | --- | --- | --- | --- | --- | --- | --- |
| chr4 | 41888616 | 0.338242 | 1/1 | 0/1 | 0/0 | 0/0 | 0.9863702 | 0.462168000 |
| chr4 | 41896891 | 0.378157 | 1/1 | 0/1 | 0/0 | 0/0 | 0.9894787 | 0.446830000 |
| chr4 | 42009953 | 0.966690 | 0/0 | 0/0 | 0/0 | 1/1 | 0.9988040 | 0.008179960 |
| chr4 | 42012245 | 0.943589 | 0/0 | 0/0 | 0/0 | 1/1 | 0.9870870 | 0.000000000 |
| chr4 | 42016927 | 0.974047 | 0/0 | 0/0 | 0/0 | 1/1 | 0.9988040 | 0.000511247 |
| chr4 | 42103725 | 0.511864 | 0/0 | 0/0 | 1/1 | 0/0 | 0.9971306 | 0.366564000 |
| chr4 | 42112654 | 0.398671 | 1/1 | ./. | 1/1 | 0/0 | 0.9679579 | 0.397750000 |
| chr4 | 42115457 | 0.234911 | 1/1 | 1/1 | 1/1 | 0/0 | 0.9732186 | 0.459100000 |
| chr4 | 42121498 | 0.635972 | 1/1 | 1/1 | 1/1 | 0/0 | 0.9600670 | 0.186605000 |

Several demographic scenarios could explain this pattern. One possibility is direct introgression from Denisovans into the ancestors of present-day modern humans. Indeed, Denisovan-like haplotypes at *SLC30A9* have previously been reported in Papuan and Melanesian populations using independent methods and interpreted as Denisovan (adaptive) introgression signals^38,9^. However, this inference relies on a limited number of derived alleles shared uniquely with the Denisovan genome and absent in Neanderthal genomes (Figure 2B).

An alternative explanation is indirect introgression via Neanderthals, which themselves experienced gene flow previously from Denisovans—a scenario documented at other loci (e.g., *MUC19*)^39^. Supporting this possibility, early modern human genomes such as Zlatý kůň and Ust’-Ishim—dated after Neanderthal admixture but before the main Denisovan admixture events in modern humans^40,41^—already carry a similar haplotype structure (Figure 2A). Under this scenario, Denisovan-derived fragments may have been introgressed into Neanderthals and subsequently transmitted to modern humans shortly thereafter. If this introgression occurred over a short timescale, there would have been limited opportunity for recombination in Neanderthals to break down and integrate the Denisovan sequence into a broader Neanderthal genomic background. As a result, the introgressed fragment could have been passed into modern humans as a relatively long Denisovan-like haplotype. Subsequent recombination in modern human populations may then have eroded the surrounding Neanderthal sequence, leaving a core region that appears predominantly Denisovan in origin.

### ABC testing

The findings above led us to formally explore different evolutionary models for the *SLC30A9* region and the methionine-to-valine substitution to explain the origin and global spread of the rs1047626 mutation, using a demographic model defined in Methods based on previous demographic inferences^14^. More specifically, we evaluated four competing scenarios involving either Denisovan (DENI model) or Neanderthal introgression (NEA model), ancestral human origin (ANC model), and complex interspecies gene flow (D2N model) (Figure 3). We discarded a Denisovan direct introgression to the out-of-Africa population given that such an event is not supported by widely adopted demographic models and the absence of inferred Denisovan ancestry on Ust’ Ishim and Zlaty Kun^15,40,41^.

**Figure 3.**
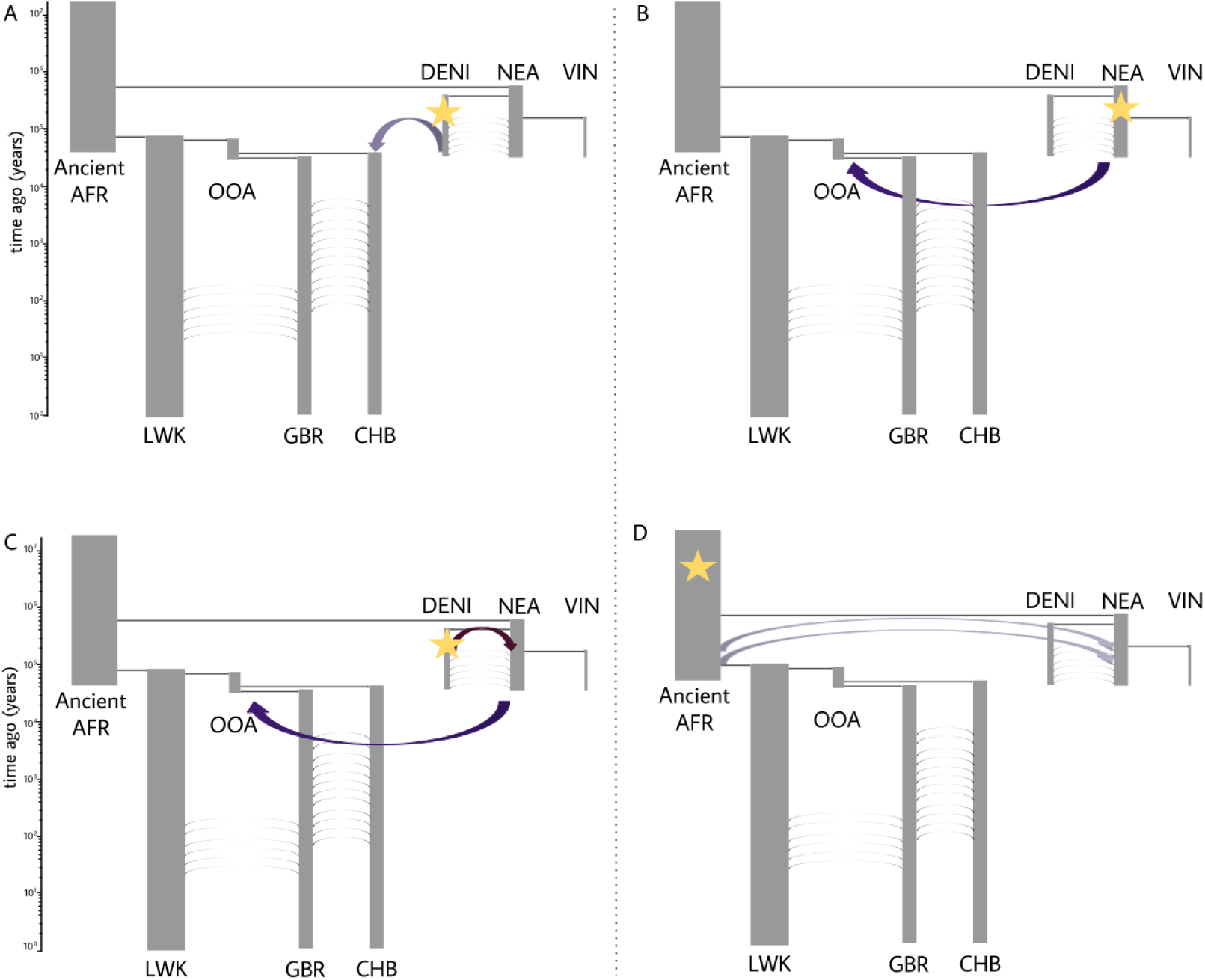
Tested evolutionary models. The star denotes the lineage where the substitution arises. In **A**, the substitution originates in Denisovans and is introgressed into East Asians. In **B**, the substitution originates in Neanderthals and is introgressed into the out-of-Africa (OOA) branch. In **C**, the substitution originates in Denisovans, introgresses first into Neanderthals, and subsequently into the OOA branch. In **D**, the substitution originates in ancient Africans and is introgressed into Neanderthals. Figure based on Jacobs 2019^14^.

We evaluated the models with ABC testing, performing forward-in-time simulations and comparing summary statistics to the empirical data derived summaries. All simulated genetic data modelled a genomic region of 310 kb using the *slendr* R package^44^ coupled with SliM^45^. As forward-in-time simulations have a big computational burden, a common practice is to scale the parameters, particularly population size and mutation rate, to reduce the number of generations. However, excessive scaling can lead to high biases in the results after a certain degree of scaling^46^, or require strong assumptions about model parameters. In this study, we used a scaling factor Q = 5 to improve computational efficiency, after verifying that such scaling did not show significant differences in the simulated genetic diversity (Supplementary Figure S11; see Methods).

From each simulation we computed 47 summary statistics from the core putative introgressed region of 70kb^10^ around rs1047626, informative about key evolutionary parameters from the models. Because large sets of statistics can introduce noise and exacerbate the curse of dimensionality—where high-dimensional spaces prevent pattern recognition—we first reduced the dimensionality via Random Forest (RF)–based feature selection using the *ranger* implementation^47^. We first trained an RF model to rank statistics by variable importance and retained the most informative subset similar to *abcrf* package approach^48^ (see Supplementary Table S3 for descriptions and the full list). This selected panel was then used in the subsequent ABC-RF analyses. For identifying the most informative summary statistics, for each model we generated 50,000 simulations, considering two-thirds of the simulations in the training of the RF model and the remaining one-third for replication. A first feature we considered as *sine qua non* in the simulated dataset was the presence of the mutation in Eurasian populations at present times. The number of simulations that passed this criterion was 3,644 for the DENI model, 2,818 for the NEA model, 4,192 for the D2N model, and 61 for the ANC model, which already suggests that some of the proposed models are more plausible than others for explaining the presence of the SNP. Within the retained simulations, the statistics identified by the RF to best discriminate between our four considered evolutionary models were: the frequency of the allele in out-of-Africa (OOA) populations, the presence of the allele in the Denisovan genome, the admixture fD statistic and the ratio sequence divergence (Rd) between CHB and Denisovans, and between EUR and Denisovans, and finally the frequency of the allele in Neanderthals (see Supplementary Table S3). Collectively, these summary statistics assess the extent of allele sharing and genetic affinity between modern human populations and archaic hominins. We next assessed the performance of the ABC framework for model selection using the selected summary statistics. The overall accuracy of the ABC-RF cross-validation error was 0.92. The lowest sensitivity was found in the classification of the ANC and D2N models (0.867 each), whereas their specificities were high (1 and 0.978, respectively). We found the lowest positive predictive value, which measures the proportion of true positive values among all the positive values, for the NEA model (0.844) (Supplementary Table S4). These results suggested that the ABC–RF approach effectively discriminated between models (Supplementary Table S4). Furthermore, we run the ABC with Neural Network (ABC-NN) method^49^ using all the computed statistics, finding that ABC-RF outperformed ABC-NN (Supplementary Tables S4, S5).

Subsequently, we generated an extra 100,000 simulations for each model considering the ascertained summary statistics by RF to be used in the ABC inference. The number of simulations that passed the criterion of the mutation being present in Eurasian populations at the end of each simulation was 7,398 for the DENI model, 5,625 for the NEA model, 8,516 for the D2N model, and 119 for the ANC model.

The ABC analysis rejected all simulations from models proposing either an African ancestral origin of the substitution (ANC) or a Denisovan origin with direct introgression into East Asians (DENI). Similarly, the model suggesting a Neanderthal origin (NEA) showed a quite reduced (0.005) posterior probability. In contrast, the model positing a Denisovan origin followed by introgression through Neanderthals (D2N) achieved the highest posterior probability (0.995). Consequently, D2N has a Bayes factor of 242 compared to NEA. Bayes factors above 30 are consistent with strong model support^50^. Therefore, these results identify D2N as the most plausible scenario for the methionine-to-valine substitution. We next tested the model adequacy of the ABC. In Bayesian analysis, model adequacy concerns whether the joint specification of likelihood and priors can plausibly generate data resembling the observations, beyond merely yielding precise estimates or outperforming alternatives^51^. Assessment was conducted by projecting in a lower-dimensional space by means of principal component analysis (PCA) the summary statistics estimated in the different simulations and the observed data, verifying that the observed statistics falls under the D2N simulations and that RF features clearly discriminates between models (Supplementary Figure S12).

We further concentrated on estimating the evolutionary parameters of the D2N model. First, for each parameter, we built an RF model to identify the most informative summary statistics. Next, we evaluated the accuracy of parameter estimation by assessing how well the posterior means obtained through ABC reflected the true values used to generate the simulated data. To this end, we treated sampled simulated datasets as observed data and re-ran the ABC inference inspecting how similar the mean of the posterior distribution of a given parameter was the one used in the simulation. To that end, we calculated the Factor 2 statistic, which measures the proportion of cases in which the true parameter value falls within 50–200% of the posterior mean estimate^52^. In our analyses, Factor 2 was more than 80% for all the inferred parameters with the exception of the selection coefficient in GBR (79%), suggesting the mean of the posterior distribution as a reliable statistic of the true model parameter. We also computed the correlation between the mean posterior distribution and the true parameter value. All parameters show significant correlation, except the selection coefficient inference in Denisovans (r^2^ = -0.024, p-value = 0.36), probably because the Denisovan population is only represented with one sample. Among the most important features extracted by the RF to infer the selection coefficient, the most important is the allele frequency of the substitution in each population, estimates of genetic diversity (the number of segregating sites and Watterson’s *θ*) and admixture statistics (Supplementary Table S5).

Given these results, which suggests that ABC was able to extract the posterior distributions of the different parameters, we performed parameter inference on the observed data using ABC-RF with local linear regression on the test simulations generated under the D2N model and the extracted features for each parameter. For all the considered parameters, we observed deviations of the posterior distributions from the prior distributions (Figure 4, Table 2). The highest median selection coefficient (s) were obtained for the Denisovan (s=0.033, 95% credible interval (CI) = [0.006 - 0.050]) and Han Chinese (s= 0.028 [0.011 - 0.048; 95% CI]), followed by the OOA populations (s=0.023, [-0.0006 - 0.048; 95% CI]) and Neanderthals (s=0.015, [-0.007 - 0.038; 95% CI]). Additionally, the appearance of the mutation in Denisovans was estimated to be 354,346 years (with a 95% CI of [280,480 - 407,741] years) (see Table 2). We generated 100,000 simulations under the D2N model using the previously inferred posterior probabilities in order to evaluate model adequacy^51^. PCA lower-dimensional projection shows that the observed statistics resemble the observed data (Supplementary Figure S13A). Moreover, we found that only 14% of the simulations have a higher distance to the center of the PCA than the observed data, suggesting that the model adequately resembles the observed genetic diversity (Supplementary Figure S13B).

**Figure 4.**
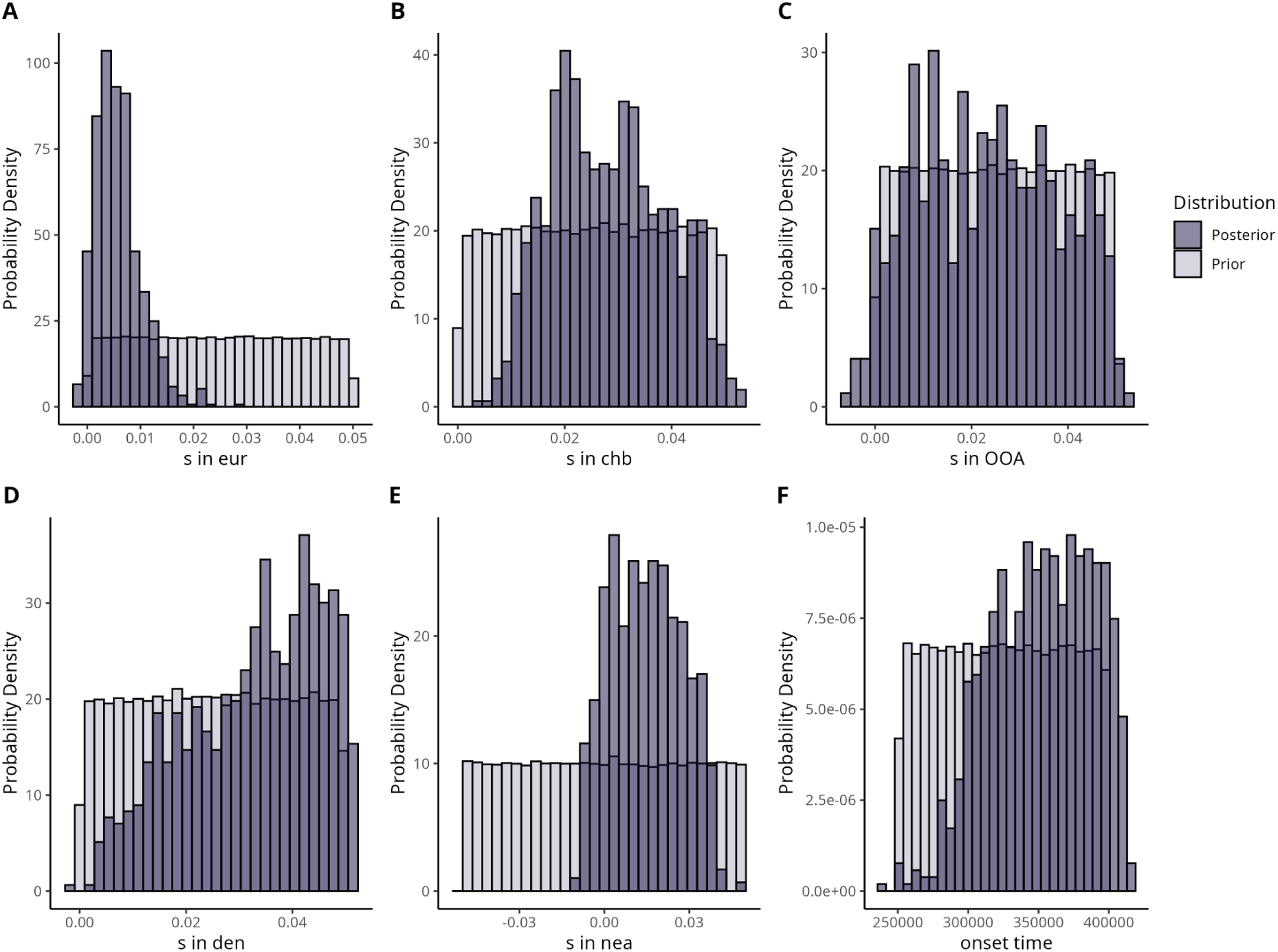
Inference of the posterior distributions (in dark purple) using ABC-RF with the local linear regression method. In light purple, the prior distribution. **A)** Posterior distribution for the selection coefficient (s) in European individuals (GBR). **B)** Posterior distribution for s in East Asian individuals (CHB). **C)** Posterior distribution for s in out-of-Africa. **D)** Posterior distribution for s in Denisovans (DEN). **E)** Posterior distribution for s in Neanderthals (NEA). **F)** Posterior distribution for the time of appearance of the mutation in Denisovans. All posterior distributions are statistically different from the priors (p-value < 0.05) after using the Kolmogorov-Smirnov (KS) test. See Table 2 for summary statistics values describing the distributions.

**Table 2.**
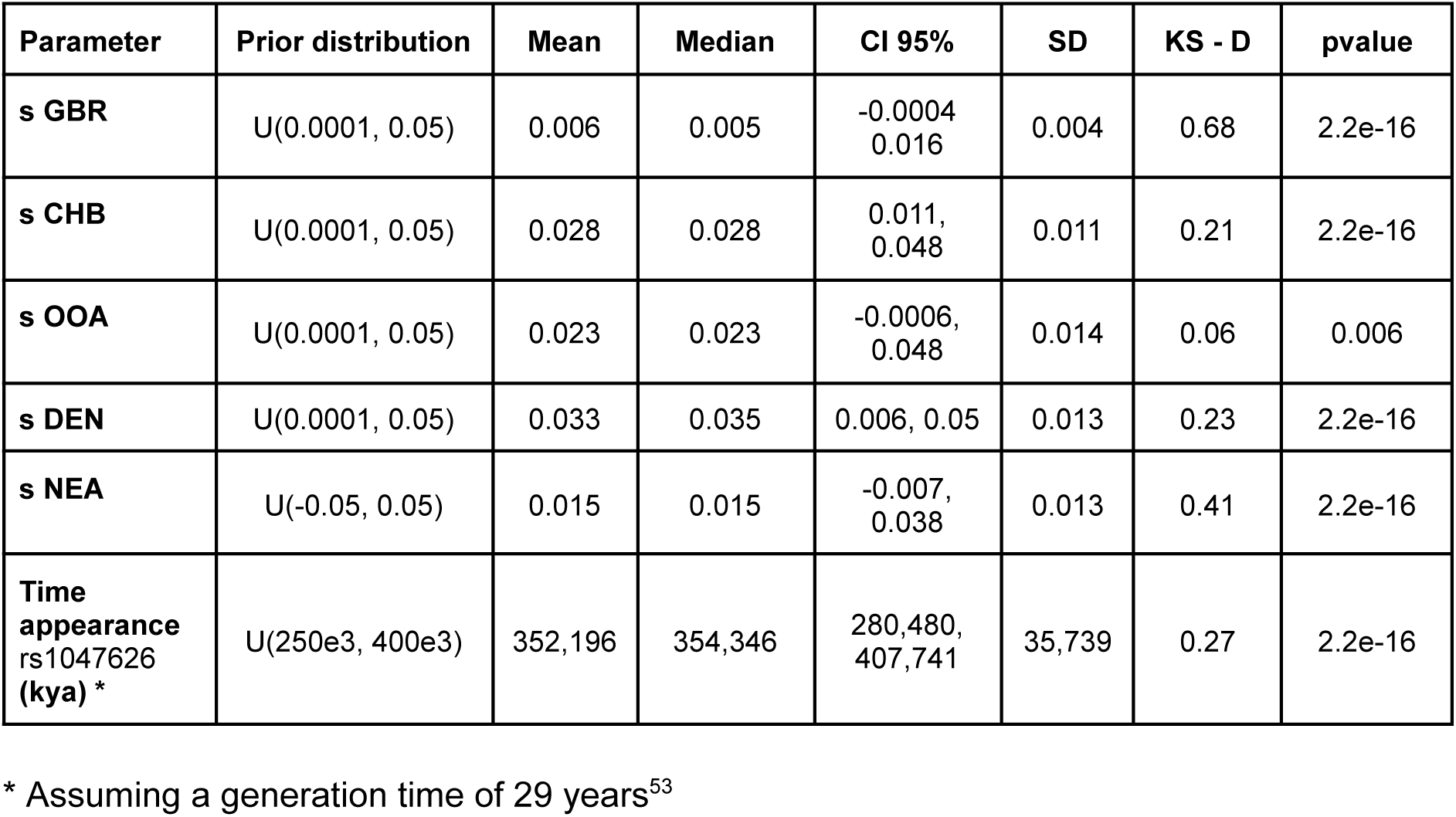
Posterior distribution statistics and prior-posterior comparisons for inferred evolutionary parameters. Table of centrality (mean, median) and dispersion (95% credible interval, standard deviation) statistics from the inferred posterior distributions for the considered parameters: selection coefficient in European individuals (s GBR), in East Asian individuals (s CHB), in the out-of-Africa population (s OOA), in Denisovans (s DEN), in Neanderthals (s NEA), and the time of appearance of the methionine-to-valine substitution. Distances between the prior and posterior distributions and p-values of the true difference between distributions calculated using the Kolmogorov-Smirnov (KS) test.

| Parameter | Prior distribution | Mean | Median | CI 95% | SD | KS - D | pvalue |
| --- | --- | --- | --- | --- | --- | --- | --- |
| **s GBR** | U(0.0001, 0.05) | 0.006 | 0.005 | -0.0004<br>0.016 | 0.004 | 0.68 | 2.2e-16 |
| **s CHB** | U(0.0001, 0.05) | 0.028 | 0.028 | 0.011,<br>0.048 | 0.011 | 0.21 | 2.2e-16 |
| **s OOA** | U(0.0001, 0.05) | 0.023 | 0.023 | -0.0006,<br>0.048 | 0.014 | 0.06 | 0.006 |
| **s DEN** | U(0.0001, 0.05) | 0.033 | 0.035 | 0.006, 0.05 | 0.013 | 0.23 | 2.2e-16 |
| **s NEA** | U(-0.05, 0.05) | 0.015 | 0.015 | -0.007,<br>0.038 | 0.013 | 0.41 | 2.2e-16 |
| **Time appearance**<br>**rs1047626**<br>**(kya) \*** | U(250e3, 400e3) | 352,196 | 354,346 | 280,480,<br>407,741 | 35,739 | 0.27 | 2.2e-16 |

\* Assuming a generation time of 29 years<sup>53</sup>

## DISCUSSION

The wide geographical distribution of the methionine-to-valine substitution in *SLC30A9*, together with consistent signals of positive selection across diverse present-day populations, underscores the putative adaptive advantage of this variant, particularly outside Africa^10,12,38^. Our analyses show that sub-Saharan African populations predominantly carry the ancestral allele, in contrast to populations outside Africa where the derived allele is common (Figure 1). In cases where the derived allele reaches appreciable frequencies within Africa, this pattern can be largely explained by back-to-Africa Eurasian admixture. Overall, these results are consistent with previous studies^10,38,9^ and support an origin of the derived allele outside Africa.

Here, we re-evaluated previous claims proposing a Denisovan origin for the predominant non-African haplotype at *SLC30A9* and for the methionine-to-valine substitution (rs1047626) under strong positive selection^10^. Consistent with earlier findings, we identify a ∼145 kb Denisova-like haplotype strongly associated with rs1047626, with a remarkably similar length across populations worldwide, potentially explained due to the haplotype being under positive selection. Within this haplotype, many variants are shared with sequenced archaic genomes (Figure 2A); however, only four closely localized variants—including rs1047626—allow us to distinguish a Denisovan from a Neanderthal origin. Given the greater statistical power to detect Neanderthal-shared variants, due to both the larger number of sequenced Neanderthal genomes and the higher divergence between the Altai Denisovan genome and the introgressing Denisovan lineages^14^, we consider it likely that at least the core of this haplotype is of Denisovan origin. Nevertheless, the limited number of informative variants constrains our ability to confidently resolve the archaic source of the full ∼145 kb haplotype. To address this limitation, we complemented haplotype-based analyses with demographic inference using ABC and machine learning approaches. An additional key observation is the presence of the derived allele and its associated haplotype in the ancient individuals Ust’-Ishim and Zlatý kůň, both of which predate the main Denisovan introgression event into non-African populations and show no genome-wide Denisovan ancestry^15,40,41^. Taken together, these observations led us to exclude a model of direct Denisovan admixture into the common ancestor of non-Africans in our ABC framework. We note, however, that such a model could potentially receive strong support if explicitly included in the model selection procedure. Indeed, previous work^54^ has shown that Denisovan ancestry in present-day populations such as Icelanders is compatible with direct admixture from Denisovans into the ancestors of non-Africans. If this were true, this haplotype could be a remnant of this first contact between Denisovans and the ancestors of non-Africans. However, this scenario of direct contact cannot be distinguished from an alternative model in which Denisovan ancestry was first introgressed into Neanderthals and subsequently transmitted to modern humans^54^, as proposed here.

Our ABC model choice supports a complex scenario involving both introgression and differential selective pressures across populations. Specifically, the best-supported model indicates a Denisovan origin of the haplotype introgressed into modern humans through the major Neanderthal admixture. This conclusion is unlikely to result from biases inherent to the ABC–RF framework or the forward-simulation strategy, as power analyses assessing the impact of the scaling factor on computational efficiency revealed no substantial differences between scaled and unscaled simulations. We also compared machine learning implementations of the ABC approach (RF vs. NN) and found that the RF algorithm achieved superior overall performance for feature selection and parameter estimation (Supplementary Tables S6–S7). Similarly, even though we employed a broad range of summary statistics previously used to detect adaptive introgression in similar evolutionary models with machine learning approaches^55^, the used summary statistics may not cover the whole spectrum of the genetic diversity that can be informative for detecting events of introgression. Nevertheless, we have shown that the considered statistics have enough power for distinguishing data from the simulated models and parameters. Another statistical artifact that could explain these results is that the selected model is the least inadequate among a set of misspecified models. All simulated models are simplified approximations of complex demographic and selective processes. The different admixture processes in Africa, Europe, and Asia, after the out-of-Africa migrations, are mostly approximated using migration rates. Incorporating more refined models that, for example, included the expansions of Bantu-speaking populations across two-thirds of the African continent^25,34,56^, two or more stems in ancient Africans^57,58^, or the Neolithic movements that shaped present-day European genetic structure^59^ would have made the simulations computationally impractical. However, model accuracy tests confirmed that the summary statistics generated under the model with the highest posterior probability closely matched the observed data, suggesting that the Denisovan to Neanderthal model is robust.

Interestingly, to the best of our knowledge, a case of adaptive introgression reported to be from the Denisovan through Neanderthal to anatomically modern humans gene flow route has been only previously reported in another case, *MUC19*^39^, only documented to be adaptive in the American continent. In the *MUC19* case, evidence indicates that each Chagyrskaya and Vindija Neanderthals carried one copy of a Denisovan-like haplotype. However, unlike the *MUC19* introgressed haplotype, none of the analyzed Neanderthal genomes (*i.e.*, Altai, Vindija, Chagyrskaya, Spi, and Goyet; Supplementary Figure S14) showed evidence of the methionine-to-valine substitution at *SLC30A9* present in the Denisovan haplotype (note that in other available Neanderthal genomes, such as Mesmaizkaya, rs1047626 is either uncovered or of insufficient sequencing quality). This absence could just reflect a sampling bias or a low frequency of the derived allele in the Neanderthal population. For example, if we assume a hypothetical allele frequency of up to 20% in Neanderthals, the probability of not observing the derived allele at rs1047626 in any of the five Neanderthal genomes currently available is up to 10.73%. Another plausible explanation is that we could not have any genome available from the actual putatively introgressed Neanderthal populations. Neanderthal migration patterns have been revealed to be more complex than previously inferred. A recent model using agent based simulations has suggested Neanderthal dispersals to Asia at 60,000-40,000 kya and 110,00−70,000 kya^60^, while a new Neanderthal mitochondrial genome from Crimea has shown long Neanderthal movements into Central Asia connecting Western Neanderthals with Altai Neanderthals^61^. Accordingly, substantial *SLC30A9* haplotype differences have been described across the Neanderthal genomes nowadays available, with Altai Neanderthal, which is 120 kya, being the most similar configuration to the Denisovan genome and the putatively introgressed haplotype in humans^10^.

Finally, the absence of the derived allele in Neanderthals could reflect the presence of negative selective pressures. Consistent with this interpretation, the posterior distribution of the selection coefficient estimated for the Neanderthal population in our ABC analyses covered weakly negative selection in the Neanderthal genomic background, although positive selection was also included in the CI. In such a scenario of negative or nearly neutral selection, the derived allele could have been passed to Neanderthals from Denisovans just very few generations before being introgressed into the Eurasians. This could also explain why the haplotype present in current populations would resemble only the Denisovan one, and not a mixture of Denisovans and Neanderthals, which would have been expected if recombination have had enough time to dilute the Denisovan haplotype within the Neanderthal ancestry, such as the observed in the case of *MUC19*^39^. Similarly, given that there is only one high-coverage Denisovan genome available at the time of this study, we note that this could be biasing our estimates for the selection coefficient in Denisovans, and that other possible scenarios may be possible in which, in both Denisovans and Neanderthals, the mutation was neutral.

Interestingly, the selection coefficients in both the ancestral out-of-Africa population and the current European populations show a posterior distribution from neutral to positive. The higher median inferred for the out of Africa population (0.023) than for the European population (0.005) could imply a relaxation of the selection as the populations that interbred with Neanderthals expanded from their possible hub in the Middle East to Europe^62^. On the contrary, the high selection coefficient in East Asian populations points out that the selective pressure continued when these populations split from the basal Eurasian population. However, given that Neanderthal genomes lack the Denisovan haplotype, the reasons why these populations with a similar geographical distribution showed such different selection forces remains unclear.

The key uncertainty about why the archaic haplotype carrying rs1047626 underwent differential selective pressures across archaic and anatomically modern humans relates to its still-unknown biological function. Inferring an adaptive potential phenotype for the methionine-to-valine substitution at the organismal level remains difficult, since disruptions in zinc homeostasis and mitochondrial activity are likely to impact many cellular functions. Indeed, whereas no significant genome-wide associations have been reported for rs1047626, a PheWAS analysis suggested several consistent phenotypic associations within the psychiatric and metabolic domains^10^. A recently published genome-phenome map in primates has shown significant correlation between *SLC30A9* and traits related to the relative sizes of cerebellum, hippocampus, striatum, and the piriform lobe, as well as other traits such as maximum life expectancy, lactancy days, basophil count, and creatine ratio in urine^63^. Nevertheless, ZnT9 knockdown, led to movement defects and dwarfism in mice, indicating a role of *SLC30A9* in the regulation of the Growth Hormone (GH)^64^ that may help to explain this complex evolutionary pattern. Finally, among the four polymorphisms exclusively shared between Denisovans and non-African populations, only rs1047626 has a documented functional impact. However, the intronic variant rs2581430 (chr4:42,012,245, GRCh38)—also private to Denisovans and anatomically modern humans—is predicted to regulate a *DCAF4L1* enhancer across three cell types: kidney capillary endothelial cells, CD14-positive monocytes, and the RCC 786-O cell line^65^.

Further functional analysis of *SLC30A9* is required to fully elucidate the exact adaptive role of the non-synonymous methionine-to-valine substitution as well as its contribution affecting late hominid evolution, present-day genetic diversity and human health.

## METHODS

### Geographical distribution of SLC30A9 alleles

To represent the geographical distribution of the ancestral and derived alleles of the *SLC30A9* methionine-to-valine substitution (rs1047626 located at positions chr4: 42,003,671 and chr4:42,001,654 in the GRCh37 and GRCh38 genome assemblies, respectively), we estimated allele frequencies for a large number of worldwide populations (n= 10,933 individuals encompassing 419 populations). Information on each population and references were included in Supplementary Table S1. In particular, we included numerous populations from across the African continent and its islands with different linguistic affiliations, geographical distributions, and lifestyles. To better visualise the distribution of allele frequencies worldwide and in Africa, we plotted the results in pie charts using an in-house R script. Since numerous populations overlap in their geographical locations in Africa, a surface map was created using the Kriging spatial interpolation method^35^. Kriging calculations and visualisation of the results were performed using Surfer software v.15 (Golden Software S.L.). Robust correlation with Winsorization method (i.e. resetting the values of outliers to the 90% percentile) was performed using R package WRS2^66^ in order to account for the effect of outlier populations with a high amount of Eurasian ancestry.

### Preparation of observed statistics

We used the 30X high coverage genomes from Altai Denisova, Altai Neanderthal, Vindija Neanderthal, and Chagyrskaya Neanderthal ancient hominids (available in: http://ftp.eva.mpg.de/neandertal/), while for the modern day sequences we used African (Luhya in Webuye or LWK), East Asian (Han Chinese or CHB), and European (Great Britain or GBR) sequences from the 1000 Genomes Project Phase 3 (1kG)^43^. For the aDNA individuals spanning the two main introgression events, we used the Allen Ancient DNA Resource (AADR) repository^36^. We processed and masked ancient and modern VCFs according to available mask maps. We lifted over to hg38 the coordinates using Picard tools^67^ and available chain files. Then, we used bedtools to extract intersecting coordinates with modern populations from 1kG. Modern and ancient DNA data were then merged using bcftools^68^. Visualization was performed using the haplostrips software^69^. The determination of the ancestral allele was performed using the Ensemble^70^ EPO ancestral consensus sequence.

### Demographic model

We generated a consensus model based on available literature and past demographic models. Most parameters were taken from Jacobs’ model^9,14^, and stated otherwise. We modeled an ancient hominid population from 650ky before the present. In our demographic model, the Neanderthal-Denisovan lineage splits from this ancestral population 600kya. The Neanderthal-Denisovan split is then modeled at 400kya, while the Vindija-Neanderthal split is modeled at 130kya. The human ancestral lineage continues until the split of the East African lineage at 90kya.

Migration rate between CHB and EUR was set to 3.14e-5 individuals each generation^14^. The percentage of admixture between Denisovan and Neanderthals was set to 0.005^71^. The percentages of introgression admixture are specific for each demographic model. The admixture percentage for the Denisovan to East Asia was set to 0.002 at 43 kya^71^ (Denisovan Introgression model). Neanderthal admixture out of Africa was set to 0.02 at 47 kya^15,72^. The first wave of human to Neanderthal has an introgression percentage of 0.05 at 200kya, whereas the second wave of human to Neanderthal is 0.005 at 100kya^73^. As for the back-to-Africa admixture, we used 0.12, which is the inferred Eurasian ancestry in Gumuz populations from Ethiopia^74^.

In this fixed demographic model, we compared different evolutionary models of the origin and evolution of the rs1047626 mutation. The first evolutionary model (DENI) considers an Altai Denisovan origin for the rs1047626 mutation, followed by adaptive introgression into East Asians and subsequent spread to Europe via migration (to explain the widespread geographical distribution of the putatively selected haplotype)^14^. The second model (NEA) suggests an Altai Neanderthal origin of the substitution with introgression into humans and Denisovans^14,71^ (due to the relatively high similarity between the Denisovan and Neanderthal haplotypes, even though no available high-coverage Neanderthal genome carries the substitution). The third model proposes an Altai Denisovan origin for the methionine-to-valine substitution at *SLC30A9*, which was later introduced into humans via Neanderthal introgression (following previous migrations from Denisovans into Neanderthals)^71^. We refer to this scenario as the Denisovan2Neanderthal model (D2N). Finally, the ANC model proposes an ancestral human origin for the methionine-to-valine substitution, which was later introgressed into European and Altai Neanderthals^73^ and subsequently passed to the Denisovan population (see Figure 3). Moreover, in each model, we account for back-to-Africa migrations to describe the presence of the derived allele and the putative introgressed haplotype in several current African populations.

The selection coefficient priors of the simulations for the Denisovan, GBR, CHB population, and the OOA population, except the DENI model (where the presence of the substitution is not modeled as the Denisovan introgression is posterior to the split of the out of Africa population) were drawn from a uniform distribution U(0.0001, 0.05). The selection coefficient prior to the simulations for the Neanderthal population was drawn from a uniform distribution U(-0.05, 0.05). The timing of the substitution was drawn from a different uniform distribution for each model: Denisovan Introgression model U(250e3, 400e3), Neanderthal Introgression model U(100e3, 600e3), and Ancestral Variation model U(100e3, 650e3). The D2N model followed the Denisovan Introgression model, with the exception that the pulse of introgression comes from Neanderthals (with the same admixture times and percentages as the Neanderthal introgression model) (Table 3).

**Table 3:** Parameters differentiating the three models. Time of appearance of the mutation (in kya) is drawn from a Uniform distribution.

|  | NEA model | DENI model | ANC model | D2N model |
| --- | --- | --- | --- | --- |
| Time of appearance of the mutation (kya)* | $U(100e3, 600e3)$ | $U(250e3, 400e3)$ | $U(100e3, 650e3)$ | $U(250e3, 400e3)$ |
| Introgression from Neanderthals into OOA | 0.02 | - | - | 0.02 |
| Introgression from Denisovans into East Asians | - | 0.002 | - | - |
| Introgression from AMH to Neanderthals | - | - | 0.05, 0.005 | - |

\* Assuming a generation time of 29 years<sup>53</sup>

We simulated a genomic chunk of 310 kbp, accounting for the three regions that are putatively different after inspecting the haplotypic structure: core putative Denisovan introgressed region (70 kbp), which is the subject of the current study, the following (70 kbp), and the preceding regions (70 kbp) with extra chunks of 50 kbp to avoid recombination sharp edges. The recombination rate used was standard 1cM/mbp and we assumed a generation time of 29 years^53^.

All simulations were run in forward with the *slendr* R package^44^ and SLiM^45^ without restarting the simulation after the mutation is lost due to genetic drift. Restarting the simulations will imply biasing the probability of observing the mutation given the parameters of the simulation, and the loss of the mutation at given selection coefficients is indeed informative for the subsequent ABC model discrimination analysis.

Selection coefficients, mutation rates, recombination rates, generations, and effective population sizes were rescaled by a rescaling factor of 5. Using 100 simulations with a fixed selection coefficient and for the DENI model, we checked that the rescaling factor does not affect the simulated genetic diversity, compared to non-rescaling and with a rescaling factor of 2 (Supplementary Figure S11). Backwards recapitation of the genealogical history of the sequence was performed using an ancestral Ne of 30000^14^ and a mutation rate of 1e-8^75^. Neutral mutations were overlaid using a mutation rate of 1e-8^75^.

### Summary statistics

Observed data statistics from the simulations and real sequences were then computed using custom scripts and the Scikit-Allele package to calculate the ZnS statistic^76^. The statistics computed have been previously used in other methods to detect complex patterns of adaptive introgression and account for several aspects of the evolutionary history of DNA sequences^55^: Watterson’s *θ*_w_ ^77^, *θ*_π_ ^78^, number of segregating sites, ZnS^76^, Garud’s H ^79^, D and fD statistics (which is an unbiased version of the D or ABBA-BABA statistic)^80,81^, divergence ratio between sequences (Rd), and U(10,20,100)^7^. The frequency of the putatively selected substitution rs1047626 was also used as a summary statistic. We used the final frequency in present-day populations and the frequency in extinct populations in the time transects of the available aDNA samples^36^ (Supplementary Table S3). ABC analysis was performed using the abc package^82^, while Random Forests were run using the ranger package^47^. We computed statistics for the central core region putatively subject to introgression. PCA was calculated using the *prcomp* function R^83^. To avoid bias in the nucleotide sequence diversity, mostly affecting the number of segregating sites of the observed data, we computed the nucleotide diversity statistics using only the current-day populations’ VCFs before the intersection with the ancient DNA.

### A novel archaic haplotype painting approach

To assign the archaic haplotype present in the modern human population to either Neanderthals, Denisovans or humans; we developed a deterministic painting algorithm based on IBD sharing. Consistent with classical IBD theory, the expected length of shared segments is inversely related to the time to the most recent common ancestor, such that longer shared fragments indicate more recent genealogical relationships^84^.

For each genomic position in the target haplotype, we estimated the local extent of the shared haplotype by measuring the distance to the nearest flanking markers at which the proxy haplotype differs from the target haplotype. We therefore obtain, at each position, a local measure of the length of the IBD-like fragment between the target haplotype and each proxy haplotype. Under the assumption that, at a given position, the proxy haplotype with the longest shared fragment is the most likely carrier of the same ancestral segment, we defined the ancestry of the target haplotype at that position as the proxy population showing the largest shared region overlapping that site. In cases where several proxy haplotypes displayed identical maximal fragment lengths, the position was assigned proportionally to all such proxies. Because multiple San haplotypes were included, their shared fragment lengths were first collapsed by taking the maximum value across all San haplotypes, yielding a single San proxy score per position. For each position, proxy populations achieving the maximum shared fragment length were assigned a value of one. In the presence of ties, multiple proxy populations were retained. The assignment vector at each position was then normalized so that values summed to one, producing fractional assignments when ties occurred.

This approach is conceptually related to methods for detecting IBD segments between individuals, which rely on identifying long stretches of matching haplotypes as evidence of recent shared ancestry^84–86^. Our approach resembles conceptually related to previously proposed methods for local ancestry inference and local ancestry painting, which aim to assign genomic segments to ancestral source populations along the genome^4,87,88^. However, whereas most existing local ancestry methods rely on probabilistic hidden Markov models and allele-frequency or haplotype-emission models learned from contemporary reference populations, our framework is based on a direct, position-specific comparison of haplotype similarity between a target haplotype and a set of proxy haplotypes, and uses the local extent of shared haplotypic fragments as a proxy for identity by descent. In this sense, our method can be viewed as a deterministic, IBD-inspired local ancestry painting strategy specifically designed for archaic introgression analyses, where reference panels are sparse, highly diverged, and often represented by single high-coverage genomes rather than population samples.

We applied the algorithm to the phased haplotypes from the phased gnomAD-merged 1000 Genomes Project (1000G) and Human Genome Diversity Project (HGDP) dataset^37^. The algorithm was applied independently to haplotypes from four present-day populations: Melanesian (n=25), Papuan (n=32), CEU (European, n=263), and CHB (Han Chinese, n=200). Proxy reference haplotypes were defined using African San haplotypes (n=12) -all showing the ancestral allele for rs1047626- and archaic hominin genomes, including Chagyrskaya Neanderthal, Altai Neanderthal, Vindija Neanderthal, and Denisovan individuals. The block length tested for each position was 3500 SNPs. For archaic individuals, only homozygous SNVs were considered. The number of heterozygote positions removed for each archaic genome were: 26 in Denisova, 26 in Vindija, 70 in AltaiNeanderthal, and 101 in Chagyskaya. Positions with at least one missing allele were removed from the dataset. Markers that were missing in all haplotypes across the combined target and proxy panels were removed. To avoid uninformative sites, singletons were excluded. We selected as control haplotypes the ones lacking the derived rs1047626 derived variant from CEU (n=89) and Mandenka (n=42).

For each analysed haplotype, the normalized proxy assignment matrix was visualized as a stacked bar chart along genomic coordinates. Each bar represents a genomic position and is partitioned according to the relative support for each proxy population. The focal SNV at rs1047626 was highlighted using a vertical dashed line. All analyses and visualizations were performed in R using the ggplot2^89^, dplyr^90^, and tidyr^91^ packages

### Approximate Bayesian Computation

We first generated 50,000 simulations for each model to perform Random Forest (RF) feature extraction with *ranger*^47^, both for model discrimination as well as for parameter estimation. We split these simulations into ⅔ for the training dataset to perform the RF training and ⅓ as replicates. We used a hypergrid parameter search, testing different hyperparameters (the number of trees generated, the minimum number of nodes, the number of variables to split in each node, and sampling with replacement) of the RF to fine-tune the algorithm implementation on the training data (Supplementary Tables S8). For all analysis we used the *abc* R package^82^. We tested the accuracy of the model classification using the extracted features to perform ABC multinomial logistic regression cross-validation error over the replicates. We perform ABC multinomial logistic regression model discrimination with 100,000 simulations for each of the four evolutionary models using the extracted features to elucidate the model that is closest to the observed data. By decoupling the feature selection from the ABC algorithm, we were able to estimate Bayes factors between evolutionary models. See Supplementary Figure S15 for a diagram of the performed workflow.

For parameter inference, we only used the simulations under the most probable model (D2N) with ABC local lineal regression. To assess the accuracy of the parameter inference, we calculated the factor 2, which is the proportion of times in which the estimated value is in an interval bounded by values equal to 50 and 200% of the true value, the absolute root mean squared error (RMSE), and Spearman’s correlation on the replicate data (Supplementary Tables S5). We further evaluated the ABC neural network (ABC-NN) cross-validation error with all the statistics to compare with the RF + ABC (Supplementary Tables S6 and S7)..

## DATA AVAILABILITY

Data from archaic hominins is available at http://ftp.eva.mpg.de/neandertal/. Data from worldwide populations included in the 1000 Genomes Project Phase 3 are available at https://www.internationalgenome.org/data. Joint 1000 Genomes Project Phase 3 and HGDP data is available at https://gnomad.broadinstitute.org/downloads#v3-hgdp-1kg. Genotypes from ancient human populations are available at: https://dataverse.harvard.edu/dataset.xhtml?persistentId=doi:10.7910/DVN/FFIDCW. Data from comparative modern populations are available for the Sahelian dataset (E-MTAB-8434); Khoe-San datasets (E-MTAB-1259); Mozambique and Angola dataset (E-MTAB-8450); Combined Jewish, European, Middle Eastern SNP array dataset (https://github.com/agladstein/AJ_ABC); Cameroonian datasets (A-MTAB-679 and A-MTAB-678); Sudan and South Sudan dataset (https://doi.org/10.5061/dryad.bs06h); and Malagasy dataset (EGAS00001002549). C.A.F-L. was granted data access for the Modern African reference datasets: AfricanNeo Modern datasets (EGAS50000000007; EGAS50000000008; EGAS50000000009; EGAS500000000010; and EGAS500000000011); Western African dataset (EGAS00001002078); Fulani dataset (EGAS50000000451); Sahelian datasets (EGAS00001001610 and EGAS50000000451); African Genome Variation Project (AGVP) dataset (EGAS00001000959); EUROTAST dataset, (EGAS00001002535); Western Mediterranean dataset (EGAS00001003901); Ethiopia Genome Project dataset (EGAS00001000238); Chad dataset (EGAS00001001231); Swahili dataset (EGAS00001002569); Other African groups (EGAS00001006944); and Arabian and Iranian dataset (EGAS00001003335).

## CODE AVAILABILITY

The code used to generate simulations, process observed and simulated data, and run the ABC analysis is available from: https://github.com/JorgeGarciaC/ABCDen

## ACKNOWLEDGMENTS

This work was supported by PID2023-147621NB-I00 funded by MICIU/AEI/10.13039/501100011033 and by “ERDF A way of making Europe”. C.A.F-L was supported by Ramon y Cajal Programme (RYC2024-048273-I), funded by the Spanish Ministry of Science and Innovation (MCIN/AEI /10.13039/501100011033) and the European Social Fund Plus (ESF+). We would like to thank Leonardo Iassi and Martin Kulwhilm for their inputs and comments during this work. The authors also thank the Unidad de Excelencia María de Maeztu” CEX2024-001431-M, funded by MICIU/AEI/10.13039/501100011033, and the Scientific Computing Core Facility (MELIS-UPF).

## AUTHOR INFORMATION

J.G-C., O.L. and E.B. designed the study. C.A.F-L. contributed to the analysis of rs1047626 in diverse African and global populations. O.L developed the haplotype painting analysis. J.G-C. performed the haplotype painting analysis. M.C.M performed the LD analysis. J.G-C. designed and performed all analysis involving simulation of different evolutionary models, ABC and machine learning. J.G-C., O.L. and E.B. wrote the manuscript with contributions and comments from M.C.M and C.A.F-L.

## REFERENCES

1. Prüfer, K. et al. The complete genome sequence of a Neanderthal from the Altai Mountains. Nature 505, 43–49 (2014).

2. Chen, L., Wolf, A. B., Fu, W., Li, L. & Akey, J. M. Identifying and Interpreting Apparent Neanderthal Ancestry in African Individuals. Cell 180, 677–687.e16 (2020).

3. Larena, M. et al. Philippine Ayta possess the highest level of Denisovan ancestry in the world. Current Biology 31, 4219–4230.e10 (2021).

4. Sankararaman, S. et al. The genomic landscape of Neanderthal ancestry in present-day humans. Nature 507, 354–357 (2014).

5. Sankararaman, S., Mallick, S., Patterson, N. & Reich, D. The Combined Landscape of Denisovan and Neanderthal Ancestry in Present-Day Humans. Current Biology 26, 1241–1247 (2016).

6. Dannemann, M., Andrés, A. M. & Kelso, J. Introgression of Neandertal- and Denisovan-like Haplotypes Contributes to Adaptive Variation in Human Toll-like Receptors. The American Journal of Human Genetics 98, 22–33 (2016).

7. Racimo, F., Marnetto, D. & Huerta-Sánchez, E. Signatures of Archaic Adaptive Introgression in Present-Day Human Populations. Molecular Biology and Evolution 34, 296–317 (2017).

8. Massilani, D. et al. Denisovan ancestry and population history of early East Asians. Science 370, 579–583 (2020).

9. Gower, G., Picazo, P. I., Fumagalli, M. & Racimo, F. Detecting adaptive introgression in human evolution using convolutional neural networks. eLife 10, e64669 (2021).

10. Roca-Umbert, A. et al. Human genetic adaptation related to cellular zinc homeostasis. PLOS Genetics 19, e1010950 (2023).

11. Barreiro, L. B., Laval, G., Quach, H., Patin, E. & Quintana-Murci, L. Natural selection has driven population differentiation in modern humans. Nat Genet 40, 340–345 (2008).

12. Grossman, S. R. et al. A Composite of Multiple Signals Distinguishes Causal Variants in Regions of Positive Selection. Science 327, 883–886 (2010).

13. Rees, J. S., Castellano, S. & Andrés, A. M. The Genomics of Human Local Adaptation. Trends in Genetics 36, 415–428 (2020).

14. Jacobs, G. S. et al. Multiple Deeply Divergent Denisovan Ancestries in Papuans. Cell 177, 1010–1021.e32 (2019).

15. Sümer, A. P. et al. Earliest modern human genomes constrain timing of Neanderthal admixture. Nature 638, 711–717 (2025).

16. Haber, M. et al. Chad Genetic Diversity Reveals an African History Marked by Multiple Holocene Eurasian Migrations. The American Journal of Human Genetics 99, 1316–1324 (2016).

17. Fernandes, V. et al. Genome-Wide Characterization of Arabian Peninsula Populations: Shedding Light on the History of a Fundamental Bridge between Continents. Mol Biol Evol 36, 575–586 (2019).

18. Gladstein, A. L. & Hammer, M. F. Substructured Population Growth in the Ashkenazi Jews Inferred with Approximate Bayesian Computation. Mol Biol Evol 36, 1162–1171 (2019).

19. Hernández, C. L. et al. Human Genomic Diversity Where the Mediterranean Joins the Atlantic. Mol Biol Evol 37, 1041–1055 (2020).

20. Pagani, L. et al. Tracing the Route of Modern Humans out of Africa by Using 225 Human Genome Sequences from Ethiopians and Egyptians. The American Journal of Human Genetics 96, 986–991 (2015).

21. Hollfelder, N. et al. Northeast African genomic variation shaped by the continuity of indigenous groups and Eurasian migrations. PLOS Genetics 13, e1006976 (2017).

22. Henn, B. M. et al. Genomic Ancestry of North Africans Supports Back-to-Africa Migrations. PLoS Genet 8, e1002397 (2012).

23. Hodgson, J. A., Mulligan, C. J., Al-Meeri, A. & Raaum, R. L. Early Back-to-Africa Migration into the Horn of Africa. PLOS Genetics 10, e1004393 (2014).

24. Hammarén, R., Goldstein, S. T. & Schlebusch, C. M. Eurasian back-migration into Northeast Africa was a complex and multifaceted process. PLoS ONE 18, e0290423 (2023).

25. Fortes-Lima, C. A. et al. The genetic legacy of the expansion of Bantu-speaking peoples in Africa. Nature 625, 540–547 (2024).

26. Fortes-Lima, C. et al. Demographic and Selection Histories of Populations Across the Sahel/Savannah Belt. Mol Biol Evol 39, msac209 (2022).

27. Fortes-Lima, C. A., Diallo, M. Y., Janoušek, V., Černý, V. & Schlebusch, C. M. Population history and admixture of the Fulani people from the Sahel. The American Journal of Human Genetics 112, 261–275 (2025).

28. Vicente, M. et al. Male-biased migration from East Africa introduced pastoralism into southern Africa. BMC Biology 19, 259 (2021).

29. Triska, P. et al. Extensive Admixture and Selective Pressure Across the Sahel Belt. Genome Biol Evol 7, 3484–3495 (2015).

30. Lankheet, I. et al. Wide-scale geographical analysis of genetic ancestry in the South African Coloured population. BMC Biology 23, 219 (2025).

31. Brucato, N. et al. The Comoros Show the Earliest Austronesian Gene Flow into the Swahili Corridor. The American Journal of Human Genetics 102, 58–68 (2018).

32. Pierron, D. et al. Genome-wide evidence of Austronesian–Bantu admixture and cultural reversion in a hunter-gatherer group of Madagascar. Proceedings of the National Academy of Sciences 111, 936–941 (2014).

33. Schlebusch, C. M. et al. Genomic Variation in Seven Khoe-San Groups Reveals Adaptation and Complex African History. Science 338, 374–379 (2012).

34. Patin, E. et al. Dispersals and genetic adaptation of Bantu-speaking populations in Africa and North America. Science 356, 543–546 (2017).

35. Matheron, G. Principles of geostatistics. Economic Geology 58, 1246–1266 (1963).

36. Mallick, S. et al. The Allen Ancient DNA Resource (AADR) a curated compendium of ancient human genomes. Sci Data 11, 182 (2024).

37. Koenig, Z. et al. A harmonized public resource of deeply sequenced diverse human genomes. Genome Res. 34, 796–809 (2024).

38. Hubisz, M. J., Williams, A. L. & Siepel, A. Mapping gene flow between ancient hominins through demography-aware inference of the ancestral recombination graph. PLOS Genetics 16, e1008895 (2020).

39. Villanea, F. A. et al. The MUC19 Gene: An Evolutionary History of Recurrent Introgression and Natural Selection. 2023.09.25.559202 Preprint at 10.1101/2023.09.25.559202 (2025).

40. Fu, Q. et al. The genome sequence of a 45,000-year-old modern human from western Siberia. Nature 514, 445–449 (2014).

41. Prüfer, K. et al. A genome sequence from a modern human skull over 45,000 years old from Zlatý kůň in Czechia. Nat Ecol Evol 5, 820–825 (2021).

42. Bergström, A. et al. Insights into human genetic variation and population history from 929 diverse genomes. Science 367, eaay5012 (2020).

43. Byrska-Bishop, M. et al. High-coverage whole-genome sequencing of the expanded 1000 Genomes Project cohort including 602 trios. Cell 185, 3426–3440.e19 (2022).

44. Petr, M., Haller, B. C., Ralph, P. L. & Racimo, F. *slendr*: a framework for spatio-temporal population genomic simulations on geographic landscapes. Peer Community Journal 3, (2023).

45. Haller, B. C. & Messer, P. W. SLiM 4: Multispecies Eco-Evolutionary Modeling. The American Naturalist 201, E127–E139 (2023).

46. Dabi, A. & Schrider, D. R. Population size rescaling significantly biases outcomes of forward-in-time population genetic simulations. Genetics 229, iyae180 (2025).

47. Wright, M. N. & Ziegler, A. ranger: A Fast Implementation of Random Forests for High Dimensional Data in C++ and R. Journal of Statistical Software 77, 1–17 (2017).

48. Raynal, L. et al. ABC random forests for Bayesian parameter inference. Bioinformatics 35, 1720–1728 (2019).

49. Blum, M. G. B. & François, O. Non-linear regression models for Approximate Bayesian Computation. Stat Comput 20, 63–73 (2010).

50. Jeffreys, S. H. The Theory of Probability. (Oxford University Press, London, 1961).

51. Gelman, A. et al. *Bayesian Data Analysis*. (Chapman and Hall/CRC, New York, 2013). doi:10.1201/b16018.

52. Excoffier, L., Estoup, A. & Cornuet, J.-M. Bayesian Analysis of an Admixture Model With Mutations and Arbitrarily Linked Markers. Genetics 169, 1727–1738 (2005).

53. Fenner, J. N. Cross-cultural estimation of the human generation interval for use in genetics-based population divergence studies. American Journal of Physical Anthropology 128, 415–423 (2005).

54. Skov, L. et al. The nature of Neanderthal introgression revealed by 27,566 Icelandic genomes. Nature 582, 78–83 (2020).

55. Zhang, X. et al. MaLAdapt Reveals Novel Targets of Adaptive Introgression From Neanderthals and Denisovans in Worldwide Human Populations. Mol Biol Evol 40, msad001 (2023).

56. Grollemund, R. et al. Bantu expansion shows that habitat alters the route and pace of human dispersals. Proceedings of the National Academy of Sciences 112, 13296–13301 (2015).

57. Ragsdale, A. P. et al. A weakly structured stem for human origins in Africa. Nature 617, 755–763 (2023).

58. Schlebusch, C. M. & Jakobsson, M. Tales of Human Migration, Admixture, and Selection in Africa. Annu Rev Genomics Hum Genet 19, 405–428 (2018).

59. Allentoft, M. E. et al. Population genomics of post-glacial western Eurasia. Nature 625, 301–311 (2024).

60. Coco, E. & Iovita, R. Agent-based simulations reveal the possibility of multiple rapid northern routes for the second Neanderthal dispersal from Western to Eastern Eurasia. PLOS ONE 20, e0325693 (2025).

61. Pigott, E. M. et al. A new late Neanderthal from Crimea reveals long-distance connections across Eurasia. Proceedings of the National Academy of Sciences 122, e2518974122 (2025).

62. Vallini, L. et al. The Persian plateau served as hub for Homo sapiens after the main out of Africa dispersal. Nat Commun 15, 1882 (2024).

63. Valenzuela, A. et al. A phylogenetic protein-coding genome-phenome map of complex traits across 224 primate species. 2025.09.08.674744 Preprint at 10.1101/2025.09.08.674744 (2025).

64. Ge, J., Li, H., Liang, X. & Zhou, B. SLC30A9: an evolutionarily conserved mitochondrial zinc transporter essential for mammalian early embryonic development. Cell. Mol. Life Sci. 81, 357 (2024).

65. Gschwind, A. R. et al. An encyclopedia of enhancer-gene regulatory interactions in the human genome. 2023.11.09.563812 Preprint at 10.1101/2023.11.09.563812 (2023).

66. Mair, P. & Wilcox, R. Robust statistical methods in R using the WRS2 package. Behav Res 52, 464–488 (2020).

67. Picard toolkit. Broad Institute, GitHub repository (2019).

68. Danecek, P. et al. Twelve years of SAMtools and BCFtools. GigaScience 10, giab008 (2021).

69. Marnetto, D. & Huerta-Sánchez, E. Haplostrips: revealing population structure through haplotype visualization. Methods in Ecology and Evolution 8, 1389–1392 (2017).

70. Harrison, P. W. et al. Ensembl 2024. Nucleic Acids Res 52, D891–D899 (2024).

71. Peyrégne, S., Slon, V. & Kelso, J. More than a decade of genetic research on the Denisovans. Nature Reviews Genetics 2023 1–21 (2023) doi:10.1038/s41576-023-00643-4.

72. Iasi, L. N. M. et al. Neanderthal ancestry through time: Insights from genomes of ancient and present-day humans. Science 386, eadq3010 (2024).

73. Li, L., Comi, T. J., Bierman, R. F. & Akey, J. M. Recurrent gene flow between Neanderthals and modern humans over the past 200,000 years. Science 385, eadi1768 (2024).

74. Haber, M. et al. Chad Genetic Diversity Reveals an African History Marked by Multiple Holocene Eurasian Migrations. The American Journal of Human Genetics 99, 1316–1324 (2016).

75. Michaelson, J. J. et al. Whole-Genome Sequencing in Autism Identifies Hot Spots for De Novo Germline Mutation. Cell 151, 1431–1442 (2012).

76. Kelly, J. K. A Test of Neutrality Based on Interlocus Associations. Genetics 146, 1197–1206 (1997).

77. Watterson, G. A. On the number of segregating sites in genetical models without recombination. Theoretical Population Biology 7, 256–276 (1975).

78. Nei, M. & Li, W. H. Mathematical model for studying genetic variation in terms of restriction endonucleases. Proceedings of the National Academy of Sciences 76, 5269–5273 (1979).

79. Garud, N. R., Messer, P. W., Buzbas, E. O. & Petrov, D. A. Recent Selective Sweeps in North American Drosophila melanogaster Show Signatures of Soft Sweeps. PLOS Genet. 11, e1005004 (2015).

80. Green, R. E. et al. A Draft Sequence of the Neandertal Genome. Science 328, 710–722 (2010).

81. Martin, S. H., Davey, J. W. & Jiggins, C. D. Evaluating the Use of ABBA–BABA Statistics to Locate Introgressed Loci. Mol Biol Evol 32, 244–257 (2015).

82. Csilléry, K., François, O. & Blum, M. G. B. abc: an R package for approximate Bayesian computation (ABC). Methods in Ecology and Evolution 3, 475–479 (2012).

83. R Core Team. *R: A Language and Environment for Statistical Computing*. (R Foundation for Statistical Computing, Vienna, Austria, 2013).

84. Thompson, E. A. Identity by Descent: Variation in Meiosis, Across Genomes, and in Populations. Genetics 194, 301–326 (2013).

85. Browning, B. L. & Browning, S. R. A Fast, Powerful Method for Detecting Identity by Descent. The American Journal of Human Genetics 88, 173–182 (2011).

86. Ralph, P. & Coop, G. The Geography of Recent Genetic Ancestry across Europe. PLOS Biology 11, e1001555 (2013).

87. Maples, B. K., Gravel, S., Kenny, E. E. & Bustamante, C. D. RFMix: A Discriminative Modeling Approach for Rapid and Robust Local-Ancestry Inference. The American Journal of Human Genetics 93, 278–288 (2013).

88. Browning, S. R., Browning, B. L., Zhou, Y., Tucci, S. & Akey, J. M. Analysis of Human Sequence Data Reveals Two Pulses of Archaic Denisovan Admixture. Cell 173, 53–61.e9 (2018).

89. Wickham, H. *Ggplot2: Elegant Graphics for Data Analysis*. (Springer-Verlag New York, 2016).

90. Wickham, H., François, R., Henry, L., Müller, K. & Vaughan, D. Dplyr: A Grammar of Data Manipulation. (2026).

91. Wickham, H., Vaughan, D. & Girlich, M. Tidyr: Tidy Messy Data. (2026).

