## Supplementary Materials for "Denisovan introgression left differential selection regimes in Humans and Neanderthals on the *SLC30A9* gene"

**Supplementary Table S1:** References for the populations included in Figure 1.

**Supplementary Table S2:** European ancestry used in Supplementary Figures S1 and S2.

**Supplementary Table S3:** List of statistics used.

**Supplementary Table S4:** Cross-validation error of the ABC Random Forest method for NEA, D2N, DEN, and ANC models with a tolerance value of 0.1.

**Supplementary Table S5:** Error assessment on parameter inference with ABC local linear regression with Random Forest impurity feature for the D2N model.

**Supplementary Table S6:** Cross-validation error of the ABC Neural Net method with a tolerance value of 0.1

**Supplementary Table S7:** Error assessment on parameter inference with the ABC neural net method for the D2N model.

**Supplementary Table S8:** Hyperparameters used in the RF feature extraction for the Model discrimination analysis and the posterior parameter inference.

A B

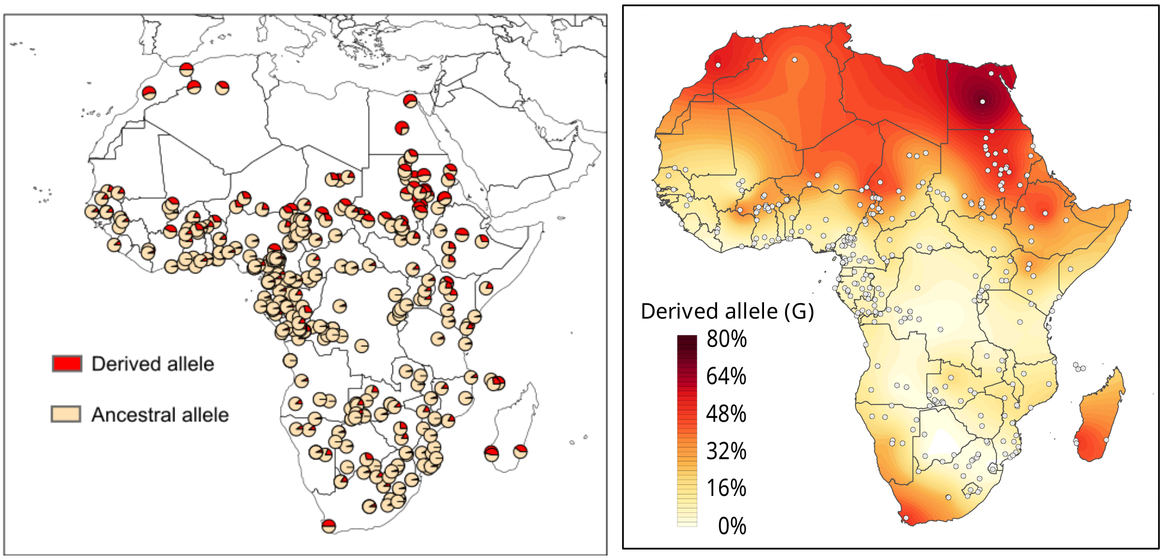


**Supplementary Figure S1:** Geographical distribution of rs1047626 allele frequencies across the African continent. **A)** Each pie graph represents the allele frequency at rs1047626 within specific modern-day African populations (see complete list and references in Supplementary Table S1). **B)** Surface map created using the Kriging spatial interpolation method. Lighter colour denotes high frequencies for the ancestral methionine variant (A allele at rs1047626), while darker colour denotes high frequencies for the derived valine variant (encoded by the G allele) of this highly differentiated nonsynonymous substitution.

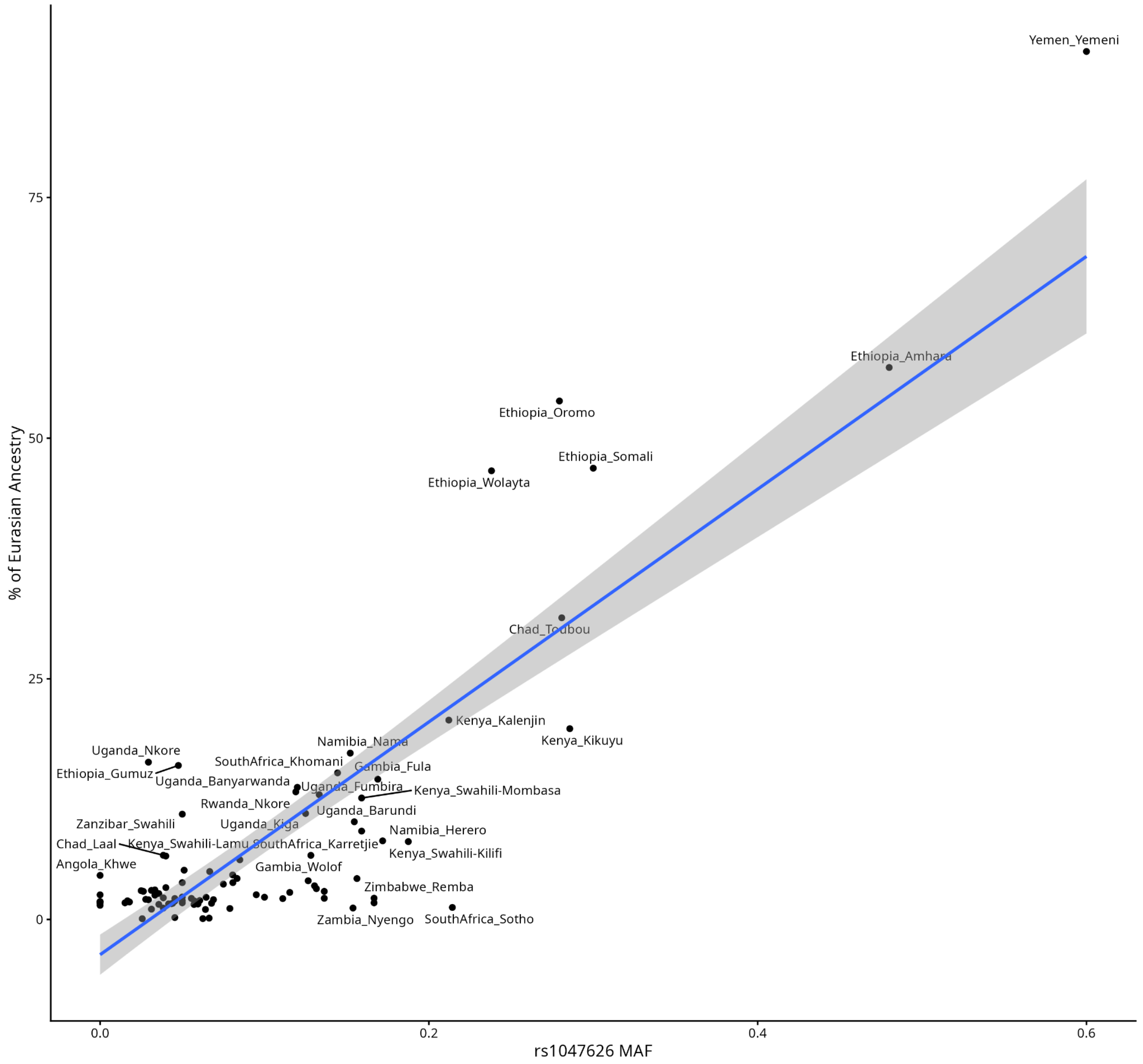


**Supplementary Figure S2:** Correlation between rs1047626 allele frequencies and Eurasian ancestry in African populations. Allele frequency at the rs1047626 locus plotted against the summed inferred percentage of European and East Asian ancestries across several African populations. Ancestry proportions were determined at K = 4 using ADMIXTURE estimates from Fortes-Lima *et al*, 2024. The blue line represents the linear model fit, with the 95% confidence interval shaded in grey. A strong positive significant correlation was observed (r = 0.5945, p-value = 1.8e-9).

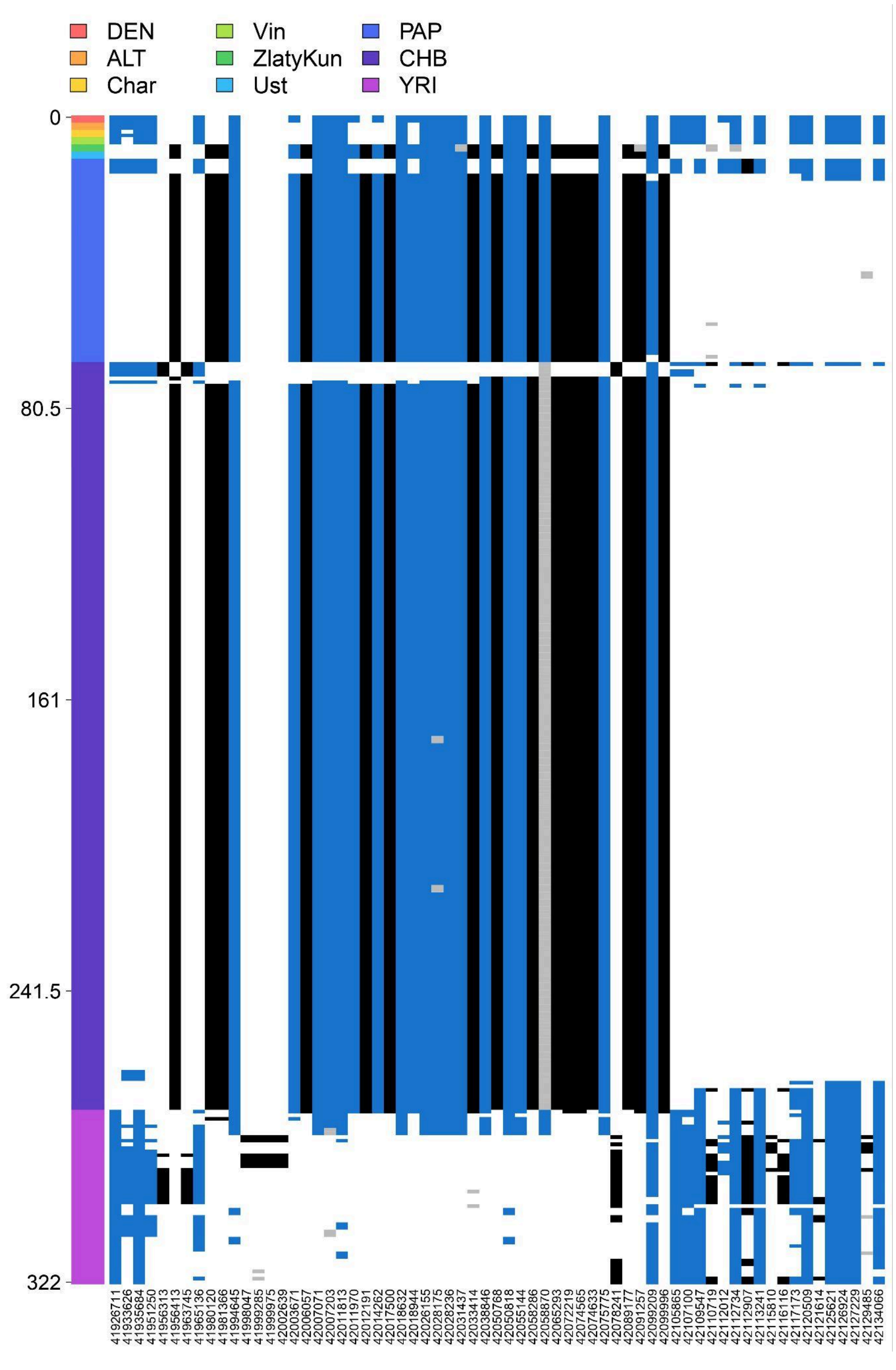


**Supplementary Figure S3:** Schematic representation of the extended haplotype structure around the *SLC30A9* gene (4:41905811-42146424, 240KB, GRCh37). In blue, derived alleles shared with Denisovans, in black, derived alleles not present in Denisovan, in white reference alleles, and in grey missing or not-called SNPs. Present-day human genomes include Yoruban (YRI) (1000 Genomes Project), Papuan (PAP) and Han Chinese (CHB) (HGDP). Archaic hominins represented are Altai Neanderthal (ALT), Altai Denisovan (DEN), Vindija Neanderthal (Vin), and Chagryskaya Neanderthal (Char). Ancient modern humans predating to 40,000 years ago include Ust'-Ishim (Ust) from Western Siberia and Zlatý kůň from the Czech Republic.

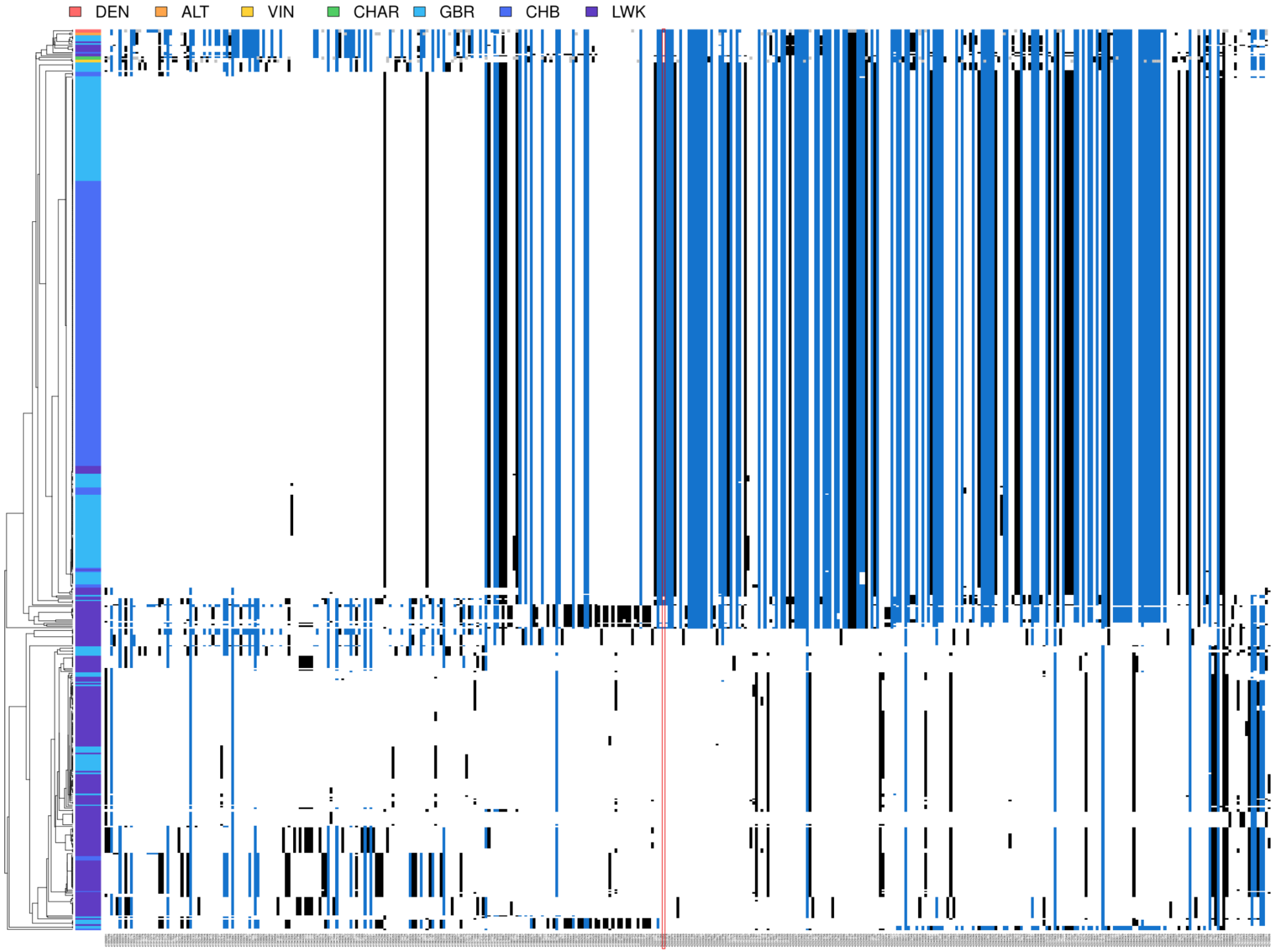


**Supplementary Figure S4:** Extended haplotype structure around the *SLC30A9* gene (4:41896891-42106606, 200KB, GRCh38). In blue, derived alleles shared with Denisovans, in black, derived alleles not present in Denisovan, in white reference alleles, and in grey missing or not-called SNPs. In red, the rs1047626 position. Present-day human genomes include Luhan in Kenya (LWK), Great Britain (GBR) and Han Chinese (CHB) from the 1000 Genomes Project. Archaic hominins represented are Altai Neanderthal (ALT), Altai Denisovan (DEN), Vindija Neanderthal (Vin), and Chagryskaya Neanderthal (Char).

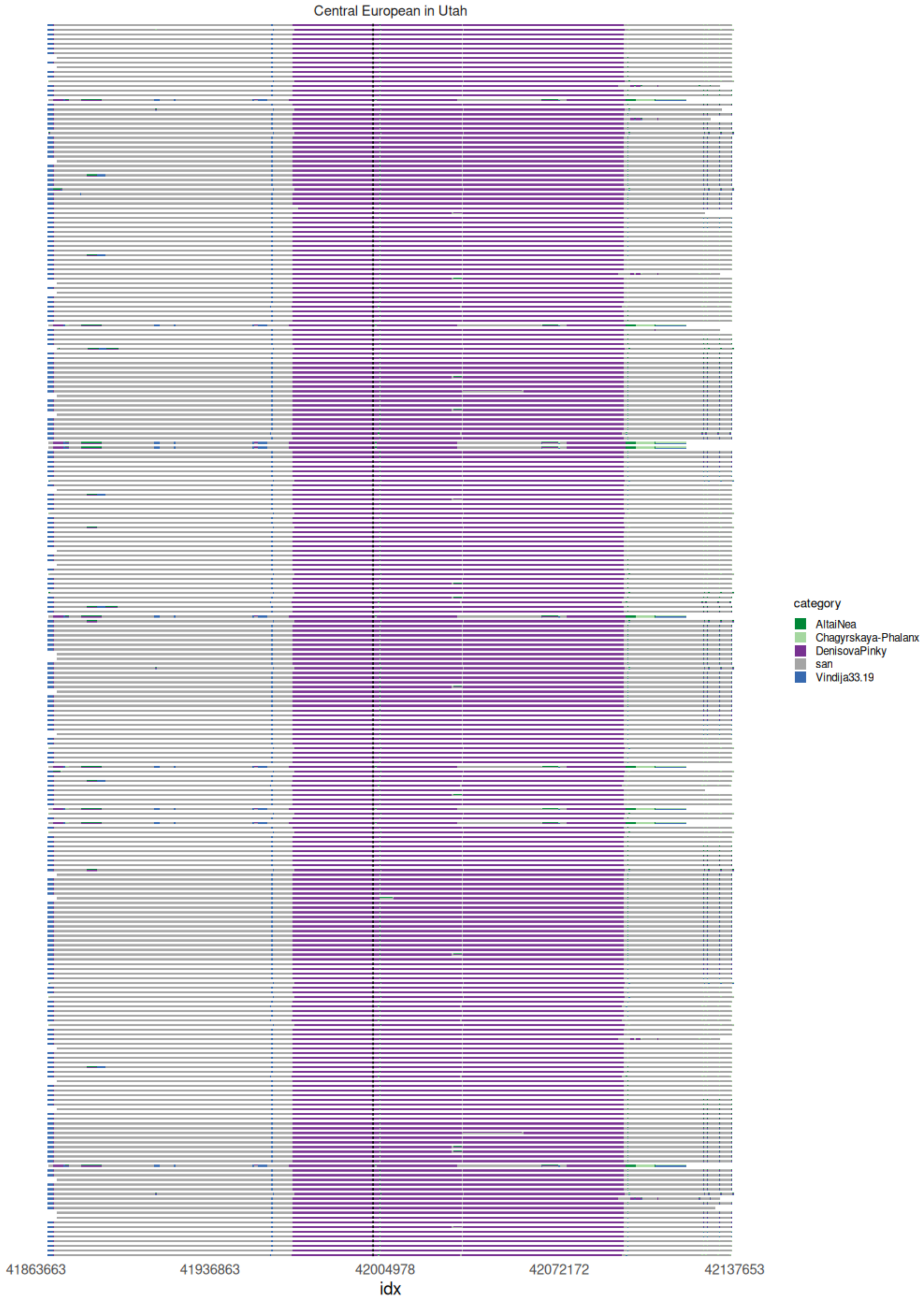


**Supplementary Figure S5:** Individual haplotypes from CEU population (n=263) painted using a set of proxy populations (San, Vindija Neanderthal, Denisovan, Altai Neanderthal and Chagyrskaya Neanderthal). White spaces denote regions not assigned to any proxy haplotype.

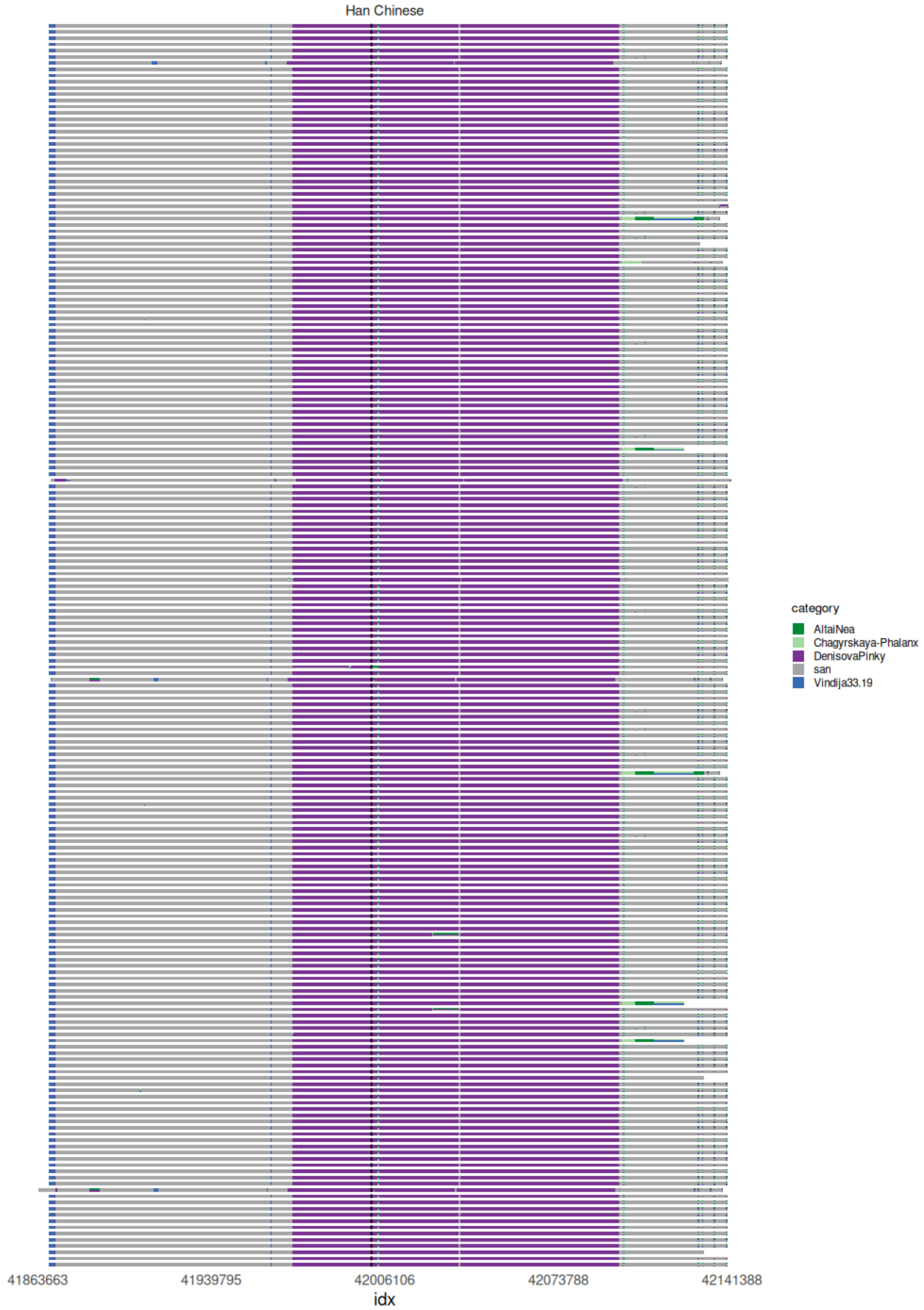


**Supplementary Figure S6:** Individual haplotypes from CHB population (n=200) painted using a set of proxy populations (San, Vindija Neanderthal, Denisovan, Altai Neanderthal and Chagyrskaya Neanderthal). White spaces denote regions not assigned to any proxy haplotype.

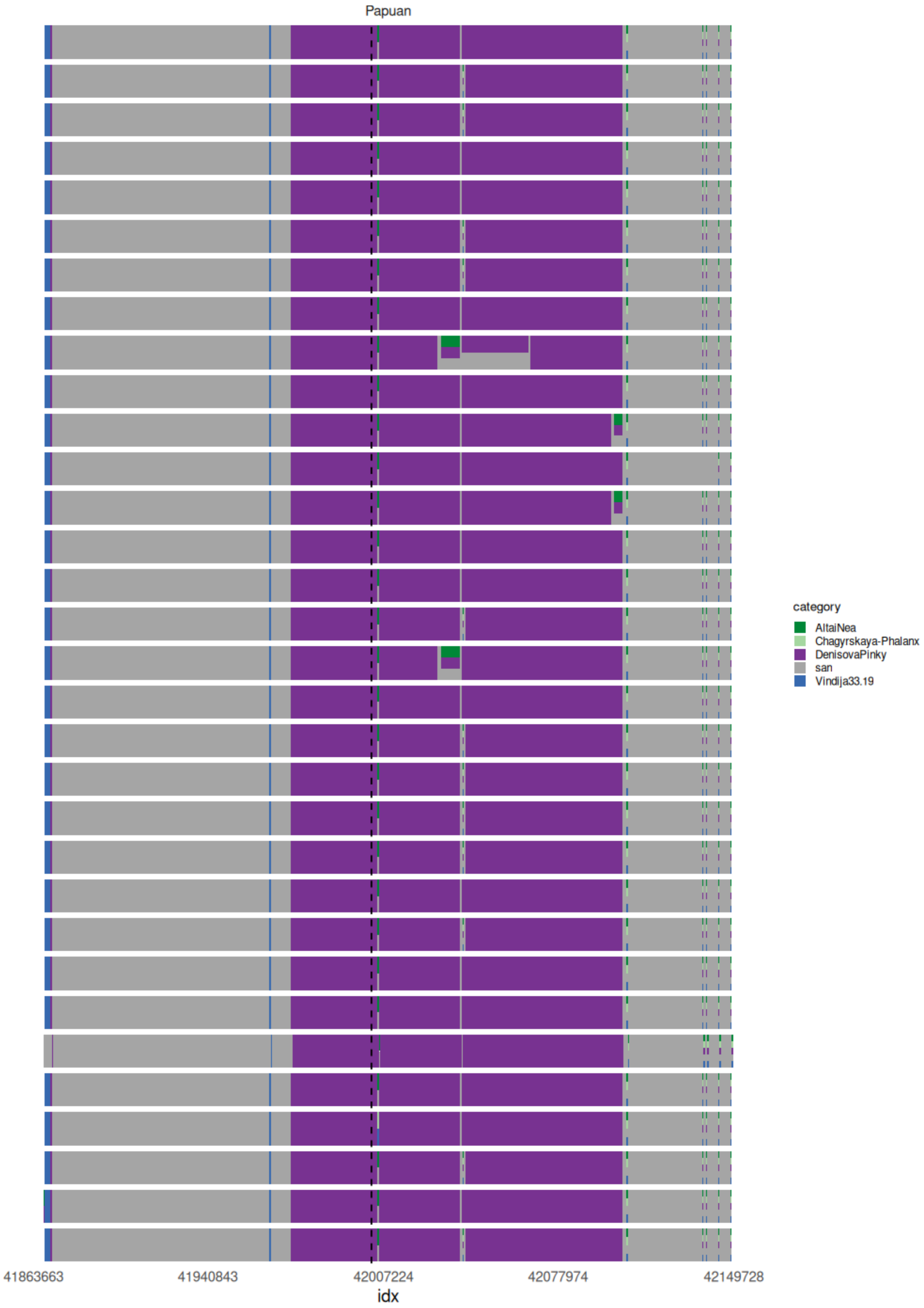


**Supplementary Figure S7:** Individual haplotypes from Papuan population (n=32) painted using a set of proxy populations (San, Vindija Neanderthal, Denisovan, Altai Neanderthal and Chagyrskaya Neanderthal). White spaces denote regions not assigned to any proxy haplotype.

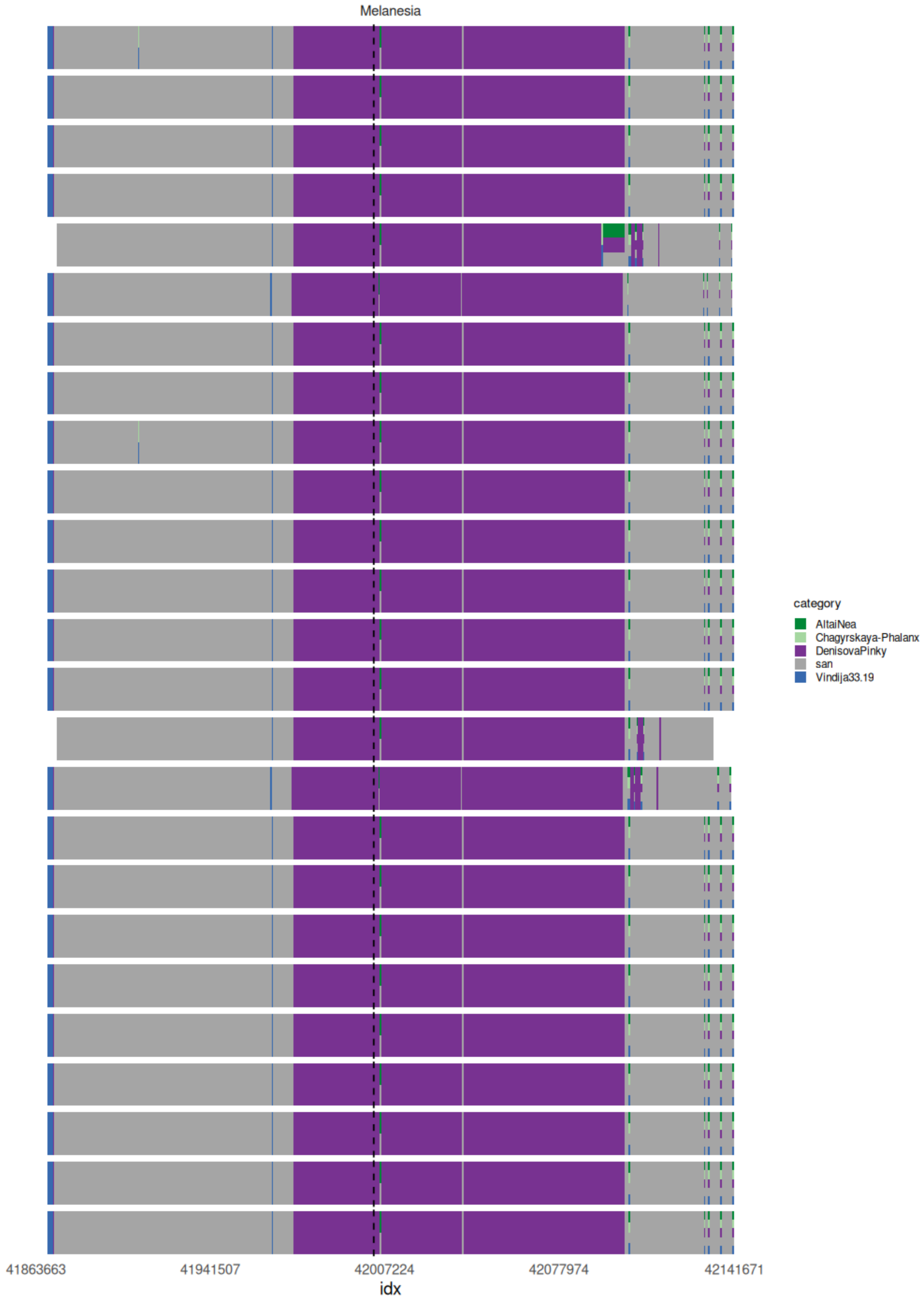


**Supplementary Figure S8:** Individual haplotypes from Melanesian population (n=25) painted using a set of proxy populations (San, Vindija Neanderthal, Denisovan, Altai Neanderthal and Chagyrskaya Neanderthal). White spaces denote regions not assigned to any proxy haplotype.

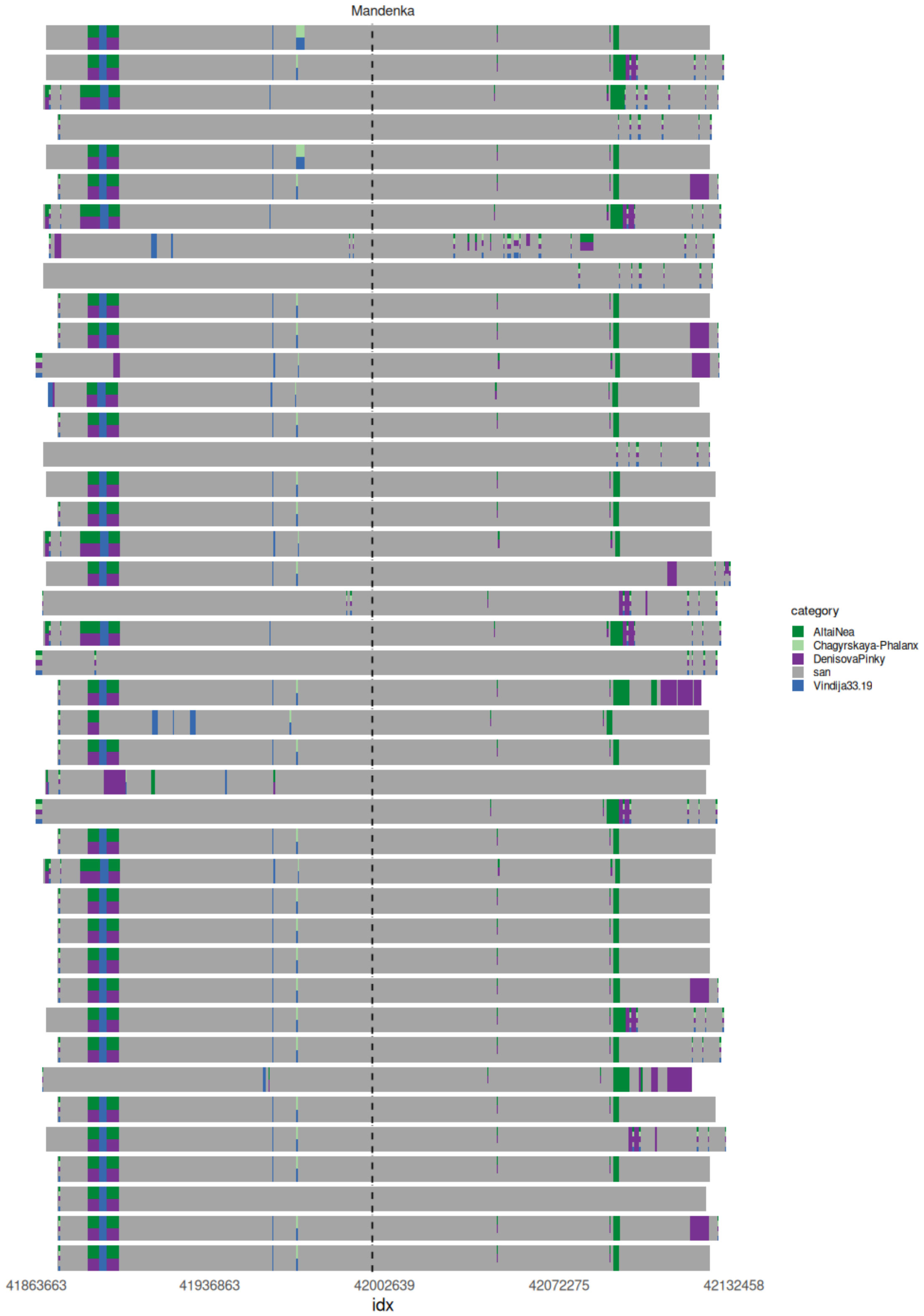


**Supplementary Figure S9:** Individual haplotypes from Mandenka population (n=42) that lack the rs1047626 derived substitution painted using a set of proxy populations (San, Vindija Neanderthal, Denisovan, Altai Neanderthal and Chagyrskaya Neanderthal). White spaces denote regions not assigned to any proxy haplotype.

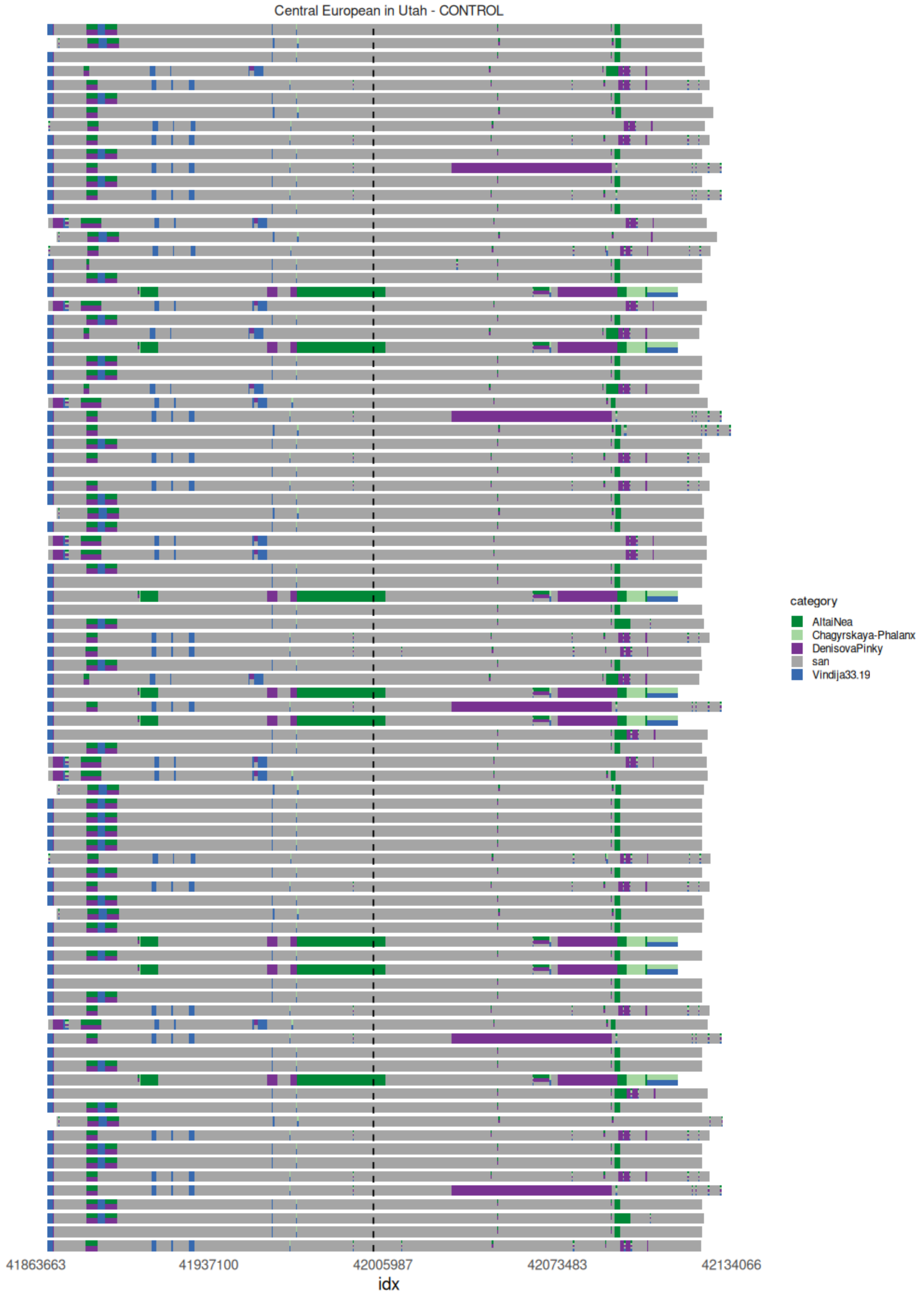


**Supplementary Figure S10**: Individual haplotypes from CEU population (n=89) that lack the rs1047626 derived substitution painted using a set of proxy populations (San, Vindija Neanderthal, Denisovan, Altai Neanderthal and Chagyrskaya Neanderthal). White spaces denote regions not assigned to any proxy haplotype.

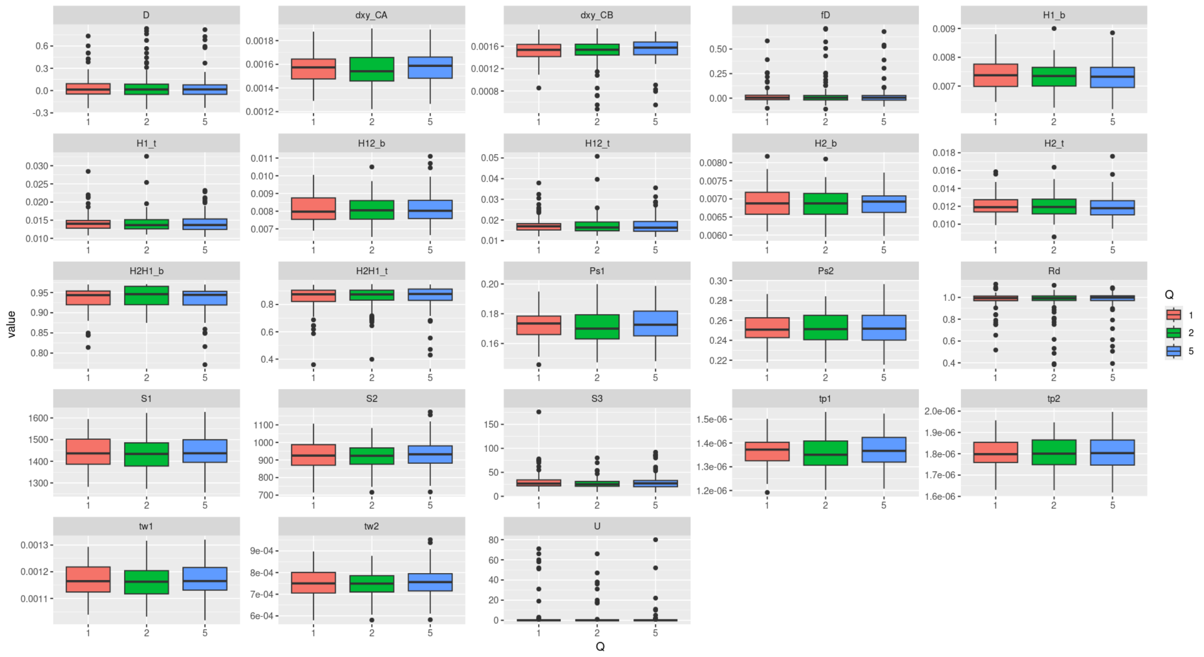


**Supplementary Figure S11:** Distribution of the computed statistics under different scaling factors (X axis). “tw” refers to the Watterson’s estimator, “dxy_CA” the divergence between Europeans and Denisovans, “dxy_CB” refers to the divergence between Europeans and African population non-admixed). “Ps” refers to the frequency of the segregating sites.

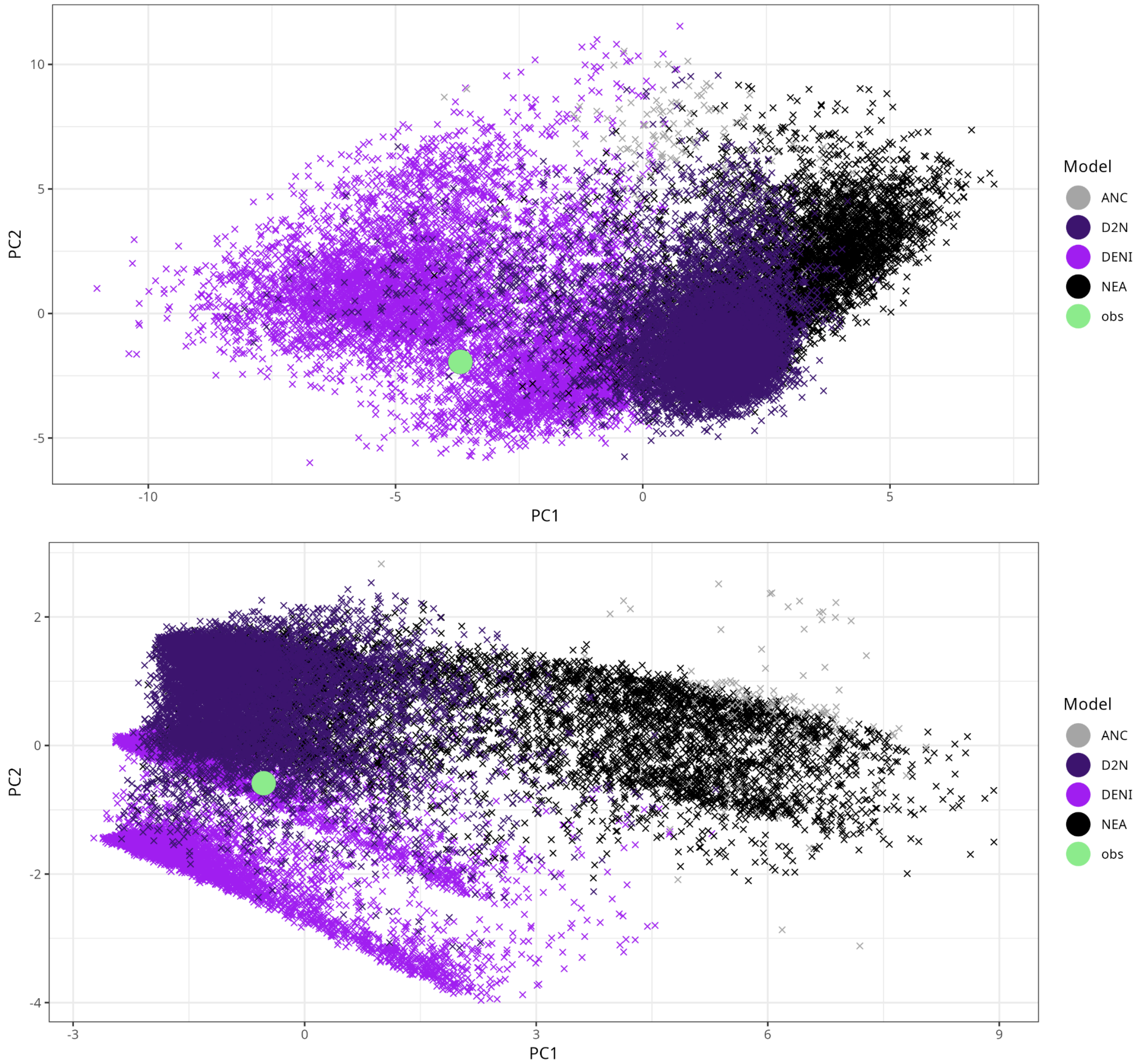


**Supplementary Figure S12:** Principal component analysis (PCA) performed with all the statistics (top) and the features that best discriminate between models using RF (bottom) for the ANC, D2N, DENI, and NEA models.

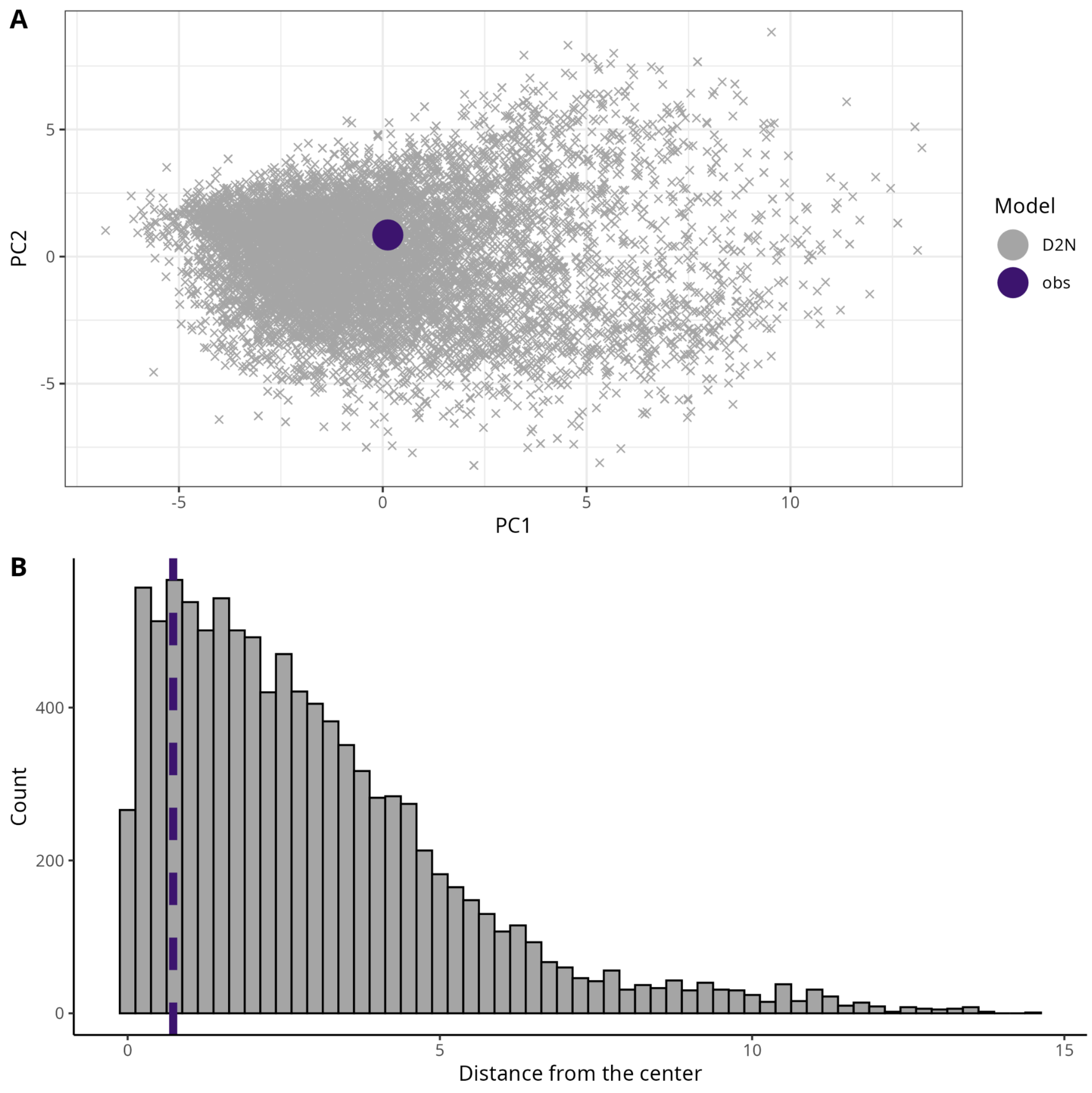


**Supplementary Figure S13: A)** PCA projection of the inferred statistics using the inferred posterior distribution under the D2N model to evaluate parameter adequacy. **B)** Distribution of distances from the first two principal components to the center. The dashed line represents the observed data, which falls near the center (p-value = 0.84).

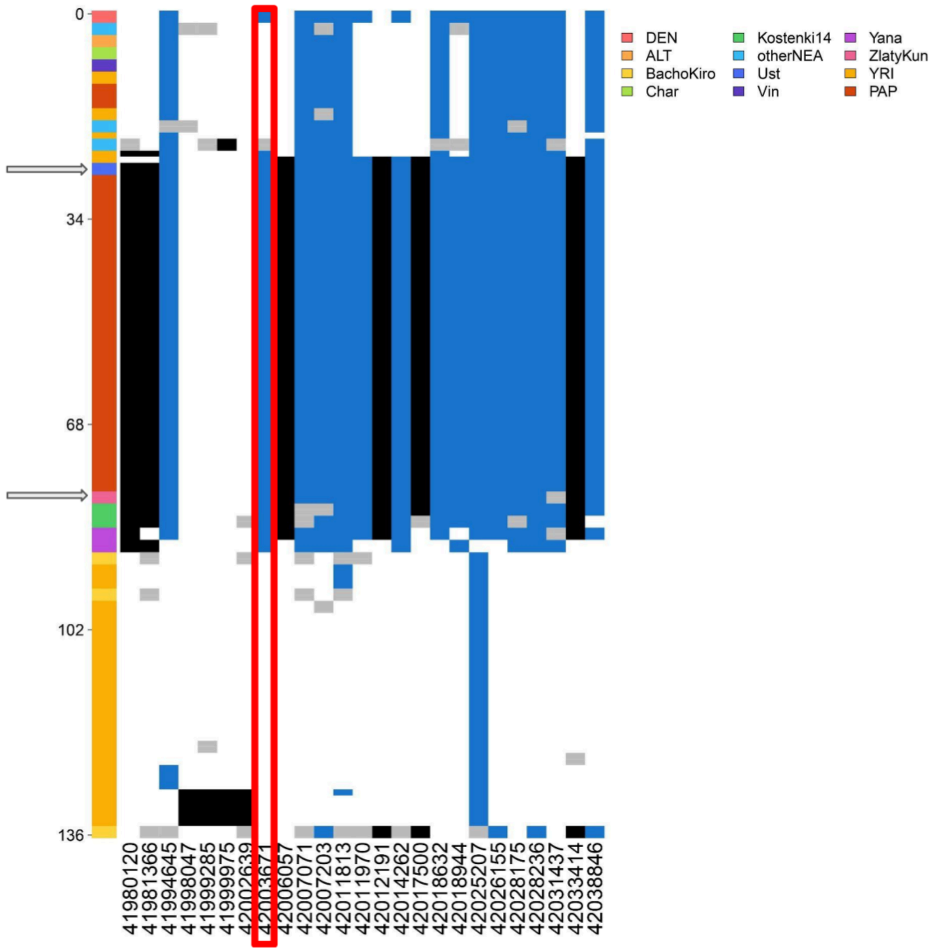


**Supplementary Figure S14:** Haplotype structure around the *SLC30A9* gene (GRCh37). As present-day genomes, we included those of the Yoruban (YRI) and Papuan (PAP) populations from the 1000 Genomes project and the HGDP, respectively. As archaic hominins, we included Altai Neanderthal (ALT), Altai Denisovan (DEN), Vindija Neanderthal (Vin), and Chagryskaya Neanderthal (Char). OtherNEA includes Goyet Neanderthal, Spi Neanderthal, and Mezmaiskaya Neanderthal. As for ancient modern humans dating between 30,000 - 40,000 ya, we included Ust'-Ishim, Zlatý kůň, Bacho Kiro, Kostenki 14, and the Yana individuals. Individuals presenting the putatively introgressed haplotype (Ust'-Ishim and Zlatý kůň) that predate the established Denisovan introgression at around 41,000 - 43,000 kya and are marked with horizontal grey arrows.

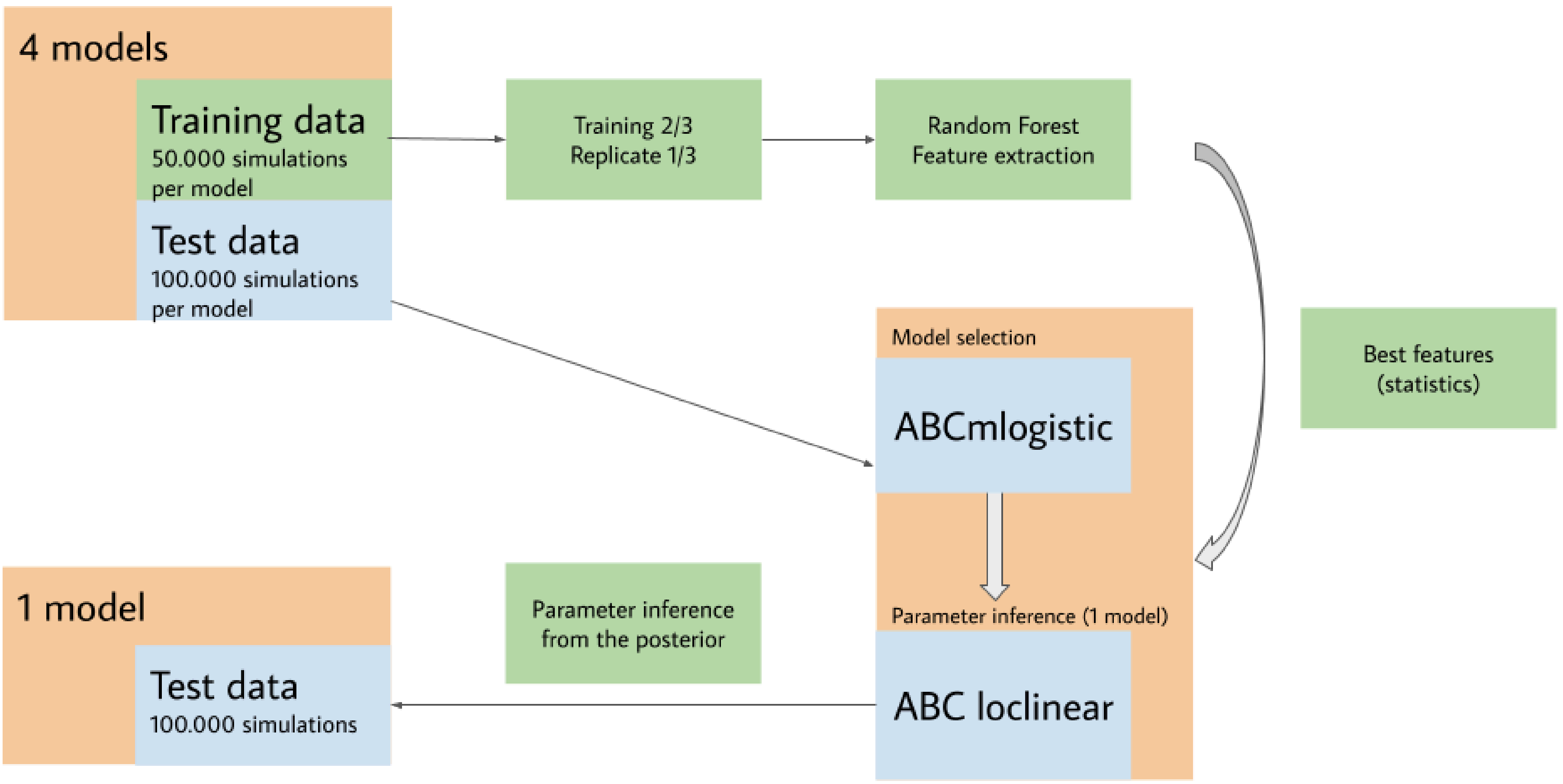


**Supplementary Figure S15:** Workflow diagram highlights the steps and the number of simulations used for the tested models.
